# Ultra-low biomass sequencing workflow (LBV-Seq) enables de novo metagenomic reconstruction of DNA and RNA viral genomes

**DOI:** 10.64898/2026.05.28.728558

**Authors:** Natalie J. Wu-Woods, Anna E. Romano, Christopher R. Neimeth, Anika Gupta, Xinyue (Penny) Pei, Adam C. Bledsoe, Joseph A. Murray, Eric V. Marietta, Rok Seon Choung, Sonia S. Kupfer, Bana Jabri, Rustem F. Ismagilov

## Abstract

Genome-resolved virome analysis remains inaccessible for many samples, including those with clinical relevance, because viral nucleic acid recovered after enrichment is often too scarce to support *de novo* genome assembly. As a result, many analyses are limited to sparse read-level detection, which cannot recover divergent viruses, resolve strains, or interpret gene-level variation. Here, we developed Low Biomass Viral Sequencing (LBV-Seq), a workflow that couples low-input viral sample handling with modified primary template-directed amplification and short- or long-read sequencing to enable *de novo* reconstruction of DNA and RNA viral genomes from sub-femtogram to nanogram inputs. LBV-Seq reproducibly captures the same relative community composition, amplifies diverse viruses, and achieved broad genome coverage across nearly all targets regardless of viral genome structure, Baltimore class, abundance, or input mass. Short-read assemblies recovered near-complete genomes from femtogram-scale inputs. Long-read sequencing provided orthogonal support for genome structure, with PacBio HiFi reads spanning large portions of viral genomes and, in some cases, complete small viral genomes. Applied to virus-enriched human duodenal biopsy eluates, LBV-Seq provided proof-of-feasibility for recovering both bacteriophage and eukaryotic viral genomes from low-input biopsy-derived material. In the eluates tested, LBV-Seq recovered co-occurring *Alphatorquevirus* and *Betatorquevirus* genomes estimated to be present at roughly 10 copies/µL and at viral masses below 0.1 fg/µL. LBV-Seq enables genome-resolved virome analysis in samples previously limited to detection-based viromics, supporting viral discovery and strain-resolved analyses in settings where viral mass is low, including viruses enriched from human tissue biopsies.

## Introduction

Viruses are major components of human-associated microbiomes and play key roles in shaping microbial communities [1], influencing host biology[2, 3], and contributing to health and disease[4, 5]. However, much of viral diversity, including in the human virome, remains poorly characterized, with a large fraction of viral sequences classified as “viral dark matter.” A key limitation is that many sample types yield insufficient viral nucleic acids for genome-resolved analyses, preventing recovery of complete viral genomes even when viral sequences are detectable [6, 7].

Genome-resolved analysis (not merely detection) is crucial. Whole-genome sequencing enables the discrimination of closely related strains and viral genes. *De novo* reconstruction enables the identification of previously uncharacterized viruses. This level of resolution is necessary to accurately infer viral function and evolutionary relationships and to robustly associate specific viral populations to biological or clinical phenotypes. However, in samples where viral nucleic acids are present at low loads, current approaches are mostly limited to sparse, read-level detection [6, 8], which indicates viral sequences are present, but cannot recover genomes absent from reference databases or resolve strain or functional variation. Targeted approaches such as probe-capture requires prior sequence knowledge and can miss divergent viruses [9].

Even when viral-enrichment methods reduce host and bacterial backgrounds, the resulting viral nucleic acid mass is often too low for robust whole-genome sequencing and de novo reconstruction [6, 10]. This limitation is particularly acute in low-biomass settings, such as aerosol samples, environment swabs, or cerebrospinal fluid. However, even high-biomass samples can yield only a small amount of viral nucleic acid mass after enrichment.

Whole-genome amplification offers one possible solution by increasing the amount of viral nucleic acids available for downstream sequencing [11]. However, the workflow must preserve the features needed for genome recovery: 1) viral community structure (relative abundances of viral taxa) must remain consistent across input biomasses 2) coverage must be distributed evenly across viral genomes, rather than concentrated in a limited number of regions, and 3) performance must remain consistent across diverse genome types and input masses. These requirements are especially challenging because viral genomes vary widely in size, strandedness, architecture, and abundance [12, 13]. Existing amplification chemistries can introduce strong biases in some viral contexts, including preferential amplification of small circular ssDNA genomes, such as with multiple displacement amplification (MDA) [14–17]. Although new methods such as primary template-directed amplification (PTA) may reduce some of these biases, they were developed largely for mammalian single-cell genomics and have not been established as workflows for genome-resolved viromics [18]. Similarly, long-read sequencing could improve viral genome reconstruction by resolving repetitive, low-complexity, or otherwise difficult-to-assemble regions, but current workflows often require more viral nucleic acid than low-biomass samples provide [19].

A key challenge is whether ultra-low-input viral material can be amplified while supporting *de novo*, reference-independent genome reconstruction from metagenomic samples. Addressing this challenge would expand the range of samples that can support genome-resolved virome analysis, including samples with low viral nucleic acid mass such as human tissues and other environments in which only small amounts of viral nucleic acid (sub-femtograms to nanograms) can be recovered.

Here, we develop and evaluate LBV-Seq (Low Biomass Viral sequencing), a sequencing workflow designed to enable de novo viral genome reconstruction from ultra-low input samples (sub-femtograms to nanograms of viral nucleic acid input mass). We optimized sample-handling and amplification for samples with low viral nucleic acid mass and evaluated performance using mock viral communities spanning genome types, abundances, and input masses. We used *de novo* assembly, rather than read mapping alone, as the primary endpoint because genome-resolved viromics requires recovery of contiguous viral genomes without relying on close reference sequences. We implemented LBV-Seq for both short- and long-read sequencing to support genome-assembly and independent assessment of genome completeness. Finally, we applied this workflow to virus-enriched human duodenal biopsies as a proof-of-principle demonstration in clinically relevant samples. Our goal was to demonstrate whether LBV-Seq workflow enables recovery of both known and divergent viral genomes from biological samples previously considered too low in viral nucleic acid input mass for *de novo* viromics.

## Results

### LBV-Seq workflow modifications enable amplification of diverse communities of low viral nucleic acid biomass

To evaluate whether the LBV-Seq workflow (Fig. 1) supports genome-resolved analysis across diverse viral genomes, we constructed DNA and RNA mock communities containing representative viruses that spanned a range of genome sizes, Baltimore classes, genome architectures, and host origins (Table 1). Each DNA and RNA mock community was evaluated across three input concentrations. DNA Mock Community 4 was serially diluted 100-fold to generate DNA Mock Communities 5 and 6, yielding a lowest total viral input mass of 150 fg in DNA Mock Community 6. RNA Mock Community 4 was serially diluted 5-fold to generate RNA Mock Communities 5 and 6, yielding a lowest total viral input mass of 360 fg in RNA Mock Community 6. Within each mock community, viruses were included at different input copy numbers so that workflow performance could be assessed across a range of relative abundances and a range of viral input masses (Fig. 2A, 2D). To quantify viral input, we designed and validated primers for qPCR assays for each virus and quantified viral copy number before amplification in each mock community. However, some communities also contained carryover host nucleic acid from incomplete virion isolation, such that total nucleic acid input mass was not always equivalent to viral nucleic acid input mass (Fig. S1).

**Table 1.** Composition of DNA and RNA mock virus communities. A mock viral community was constructed using a range of mammalian-relevant viruses and bacteriophages with varying genomic structures at differing relative abundances by mass. Viruses are ordered by genome size; relative abundances were calculated based on the mass of genomic nucleic acid input into the library preparation, as estimated by taxon-specific qPCR and genome size (see Methods Quantification qPCR).

| **DNA Mock Community 4-6** |  |  |  |  |  |
| --- | --- | --- | --- | --- | --- |
| **Virus** | **Baltimore classification** | **Host organism** | **Genome structure** | **Genome size (kb)** | **Relative abundance by mass (%)** |
| PBCV-1 | dsDNA | Chlorella (algae) | Linear | 331 | 55 |
| φKZ | dsDNA | Pseudomonas | Linear | 280 | 30 |
| Vaccinia virus | dsDNA | Homo sapiens | Linear | 195 | 9.2 |
| T4 phage | dsDNA | Escherichia | Linear | 169 | 5.3 |
| M13 phage | ssDNA | Escherichia | Circular | 6.4 | $1.3 \times 10^{-1}$ |
| AAV | ssDNA | Homo sapiens | Linear | 3.3 | $1.3 \times 10^{-3}$ |
| **RNA Mock Community 4-6** |  |  |  |  |  |
| Murine hepatitis virus | ssRNA(+) | Mus musculus | Linear | 31 | 13 |
| φ6 | dsRNA | Pseudomonas | Segmented (three segments) | 13 | 2.8 |
| Lentivirus | ssRNA(+) | Homo sapiens | Linear | 11 | 2.9 |
| Qβ | ssRNA(+) | Escherichia | Linear | 4.2 | 18 |
| MS2 phage | ssRNA(+) | Escherichia | Linear | 3.6 | 63 |

**Figure 1:**
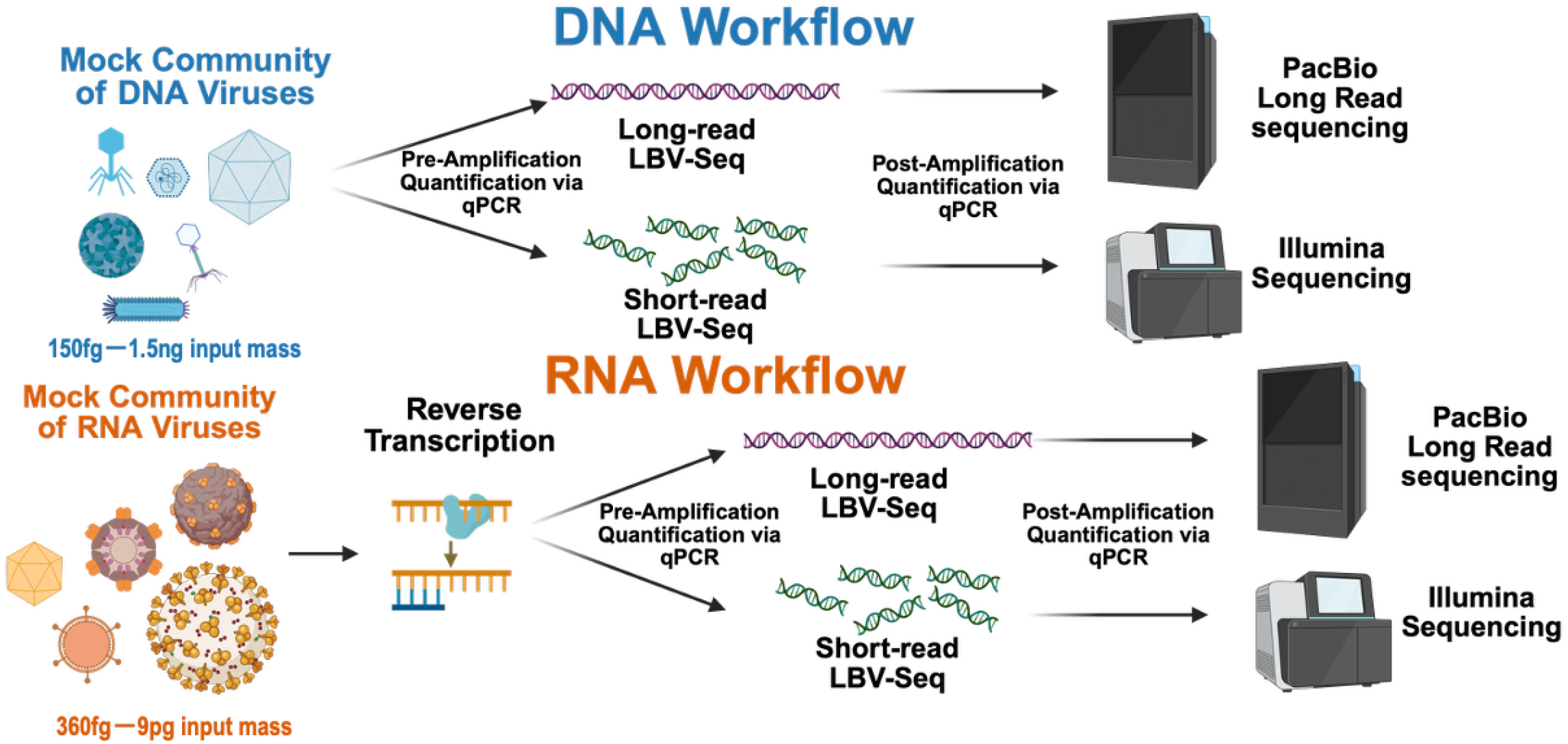
Low biomass viral sequencing (LBV-Seq) workflows for de novo viral metagenomics. Schematic of the LBV-Seq workflow developed in this study. The workflow is designed to enable genome-resolved analysis of DNA and RNA viruses from ultra-low-input samples, including human tissue.

**Figure 2:**
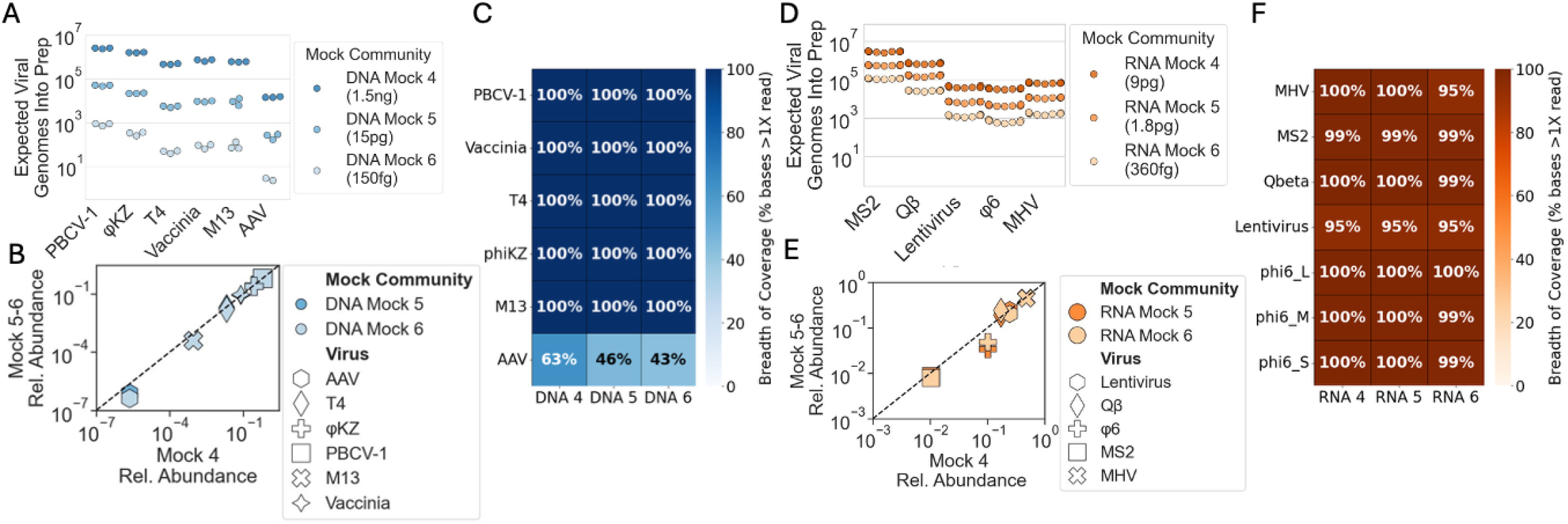
LBV-Seq workflow captures the same viral community composition and supports broad genome coverage across viral nucleic acid input masses. **A**,**D)** Expected viral genome copies entering sample preparation are shown for the DNA and RNA mock communities. DNA inputs were quantified by triplicate qPCR measurements of the same input community. RNA inputs were quantified from three reverse-transcription replicates, each measured by triplicate qPCR. **(B**,**E)** Viral relative abundances after amplification and short-read Illumina sequencing were similar across the tested input range for both the DNA (B) and RNA (E) mock communities, indicating that amplification largely captured the same community composition. **(C**,**F)** Average breadth of genome coverage for DNA viruses (C) and RNA viruses (F) after sequencing. For B, C, E, and F, each input mass was analyzed with three sequenced LBV-Seq workflow replicates. Plotted values show the mean of these three workflow replicates. Most viruses showed broad coverage across their genomes, indicating that the workflow generally supported whole-genome sequencing rather than sparse read-level detection.

We next evaluated amplification methods to select an approach that would minimize bias across viral genome types. The two main methods in the literature are MDA and PTA. MDA has non-uniform performance across genomes and overamplification of ssDNA circular viruses, whereas PTA has uniform amplification across the genome in single-cell bacteria [20] and eukaryotes [18]. To our knowledge, PTA had not been modified to work with low-biomass metagenomic viral samples. We modified the input preparation workflow so that purified gDNA or cDNA, rather than intact mammalian cells, was used as input. This modification consisted of performing upstream chemical and mechanical lysis followed by nucleic-acid purification before amplification. We also shortened and cooled the alkaline denaturation step, originally needed to lyse whole-cell inputs, to 10 min at 4 °C to minimize potential damage to short viral genomes, consistent with prior work showing that alkaline treatment can damage short DNA and cDNA molecules [21]. In addition, we increased the nucleic acid input volume from 1 µL to 3 µL to increase the amount of template entering each reaction.

Samples with low levels of viral nucleic acid (sub-femtograms) are susceptible to background contamination, so next we monitored amplification in real time with SYBR fluorescence and compared biological samples with no-template controls. We also extended the amplification time from 10 h to 14 h to maximize product yield while preserving separation from no-template-control amplification and generating sufficient material for downstream library preparation. For RNA-virus samples, we also performed bacterial and mammalian rRNA depletion before cDNA synthesis to reduce host background in downstream sequencing data.

To determine whether amplification altered community structure, we quantified viral copy number before and after amplification with qPCR (Fig. 2A,D & S2). We found that fold-change in viral abundance depended on total input mass where lower-abundance communities showed greater amplification across all viruses, resulting in similar final nucleic acid masses (Fig. S2). Overall, amplification was highly reproducible across reverse-transcription (RT) replicates, technical cDNA or DNA replicates, and input biomasses, with the coefficient of variation of post-amplification viral copy number remaining below 0.13 even when viral input mass was as low as 3.3 × 10^-3^ fg.

We next asked whether amplification performance was preserved across viral nucleic acid input masses. We compared the relative abundances measured in Mock Community 4 with those measured in the lower-biomass Mock Communities 5 and 6. Relative abundances were strongly correlated across input biomasses for both the DNA and RNA communities (R = 0.99 for DNA and R = 0.98 for RNA; Fig. 2B, 2E), indicating that amplification performance was largely preserved across the tested input range and supporting comparison of viral abundances among samples processed with the same workflow.

Because whole-genome amplification methods are known to overamplify small circular ssDNA viruses, we also tested whether the presence of ssDNA viruses distorted community composition. Comparing communities with and without ssDNA viruses showed nearly identical performance (within 1.5-fold for all viruses; R= 0.99; Fig. S3), indicating that the workflow did not introduce strong bias toward these genomes under the conditions tested.

We then evaluated whether amplification extended across the full length of each viral genome by performing short-read Illumina sequencing on the amplified products and mapping reads back to the viral reference genomes. High breadth of coverage (>90%) was achieved for nearly all viruses in both the DNA and RNA mock communities (Fig. 2C, 2F), indicating that amplification generally supported whole-genome sequencing rather than sparse detection. The main exception was AAV, which showed lower breadth of coverage, likely reflecting its extremely low relative abundance (1.3 × 10^-5^), which made recovery read-depth limited (average of 43 reads aligned to the reference). At this low target abundance, AAV recovery was likely limited by stochastic sampling of very few input molecules and by sequencing depth, so incomplete AAV coverage is consistent with Poisson dropout rather than a virus-specific failure of LBV-Seq, especially at the lower input communities. Heterogeneity in the virion preparation, for which full-length genome encapsulation is not guaranteed, has also been reported for AAV constructs [19, 22], which is another factor that may contribute to incomplete AAV genome coverage under the conditions tested. For all other viruses, coverage was broadly distributed across genomes (Fig. S4, Table S1). In a high-input dsDNA mock community that could be sequenced without whole-genome amplification, LBV-Seq preserved viral relative abundance and produced coverage profiles similar to standard workflows, such as Illumina DNA Prep (Fig. S5). This control supports that LBV-Seq enabled genome-wide recovery without introducing major additional distortions in dsDNA viral community composition or coverage.

Overall, these results demonstrate that the LBV-Seq workflow reproducibly amplifies diverse DNA and RNA viral genomes across four-order-of-magnitude input range, from femtogram to nanograms of viral nucleic acid, while preserving community structure and enabling genome-resolved analysis.

### LBV-Seq generates PacBio-compatible products and enables single-read recovery of small viral genomes

To enable long-read sequencing of ultra-low-biomass viral communities, we validated a custom long-read PTA workflow performed by BioSkryb designed to generate PacBio-compatible products from femtograms to nanograms of viral nucleic acid input mass (Fig. 1). Amplification generated sufficient material for PacBio library-preparation across the tested input range. For the DNA mock communities, amplification increased viral DNA mass by 370-fold in Mock Community 4 and by up to 96,000-fold in Mock Community 6 (final total DNA masses of 1.1–1.7 µg per reaction). RNA mock communities were submitted for amplification after cDNA synthesis and yielded lower final viral nucleic acid masses (0.5 to 0.8 µg per reaction) but were still sufficient for PacBio library-preparation without additional amplification. As in the short-read workflow, no-template controls were carried through amplification for both the DNA and RNA communities.

We next asked whether the long-read workflow successfully amplified viral nucleic acids of interest. Virus-specific qPCR of pre- and post-amplification showed amplification of viruses for all input biomasses (Fig S6). To test whether performance was maintained across input biomasses, we compared Mock Community 4 with the lower-biomass Mock Communities 5 and 6. Relative abundances remained highly correlated across biomass inputs for both the DNA and RNA communities (R = 0.99 for DNA and R = 0.99 for RNA; Fig. 3A–B), indicating that amplification behavior was largely preserved across the tested input range.

**Figure 3:**
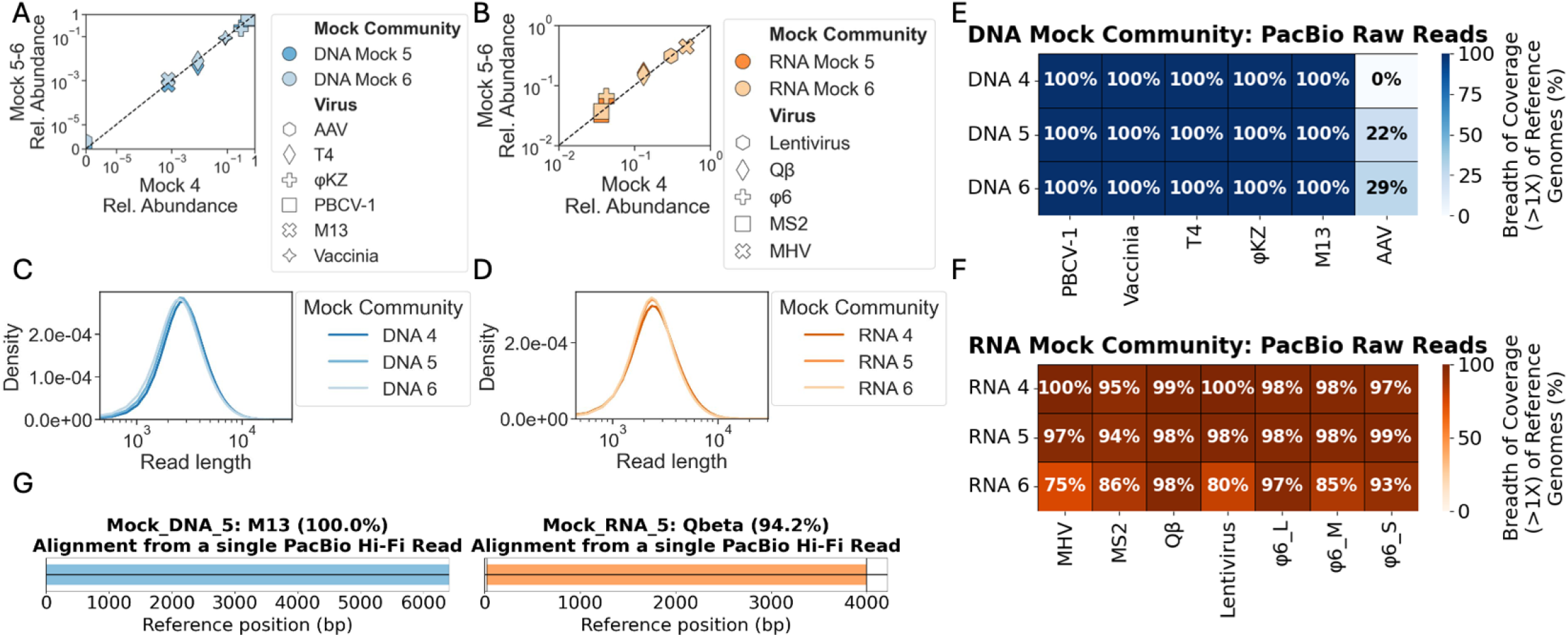
LBV-Seq for long-read generates PacBio-compatible products and supports single-read viral genome capture. **(A**,**B)** Relative abundance of each viral species after modified long-read LBV-Seq across DNA (A) and RNA (B) mock communities spanning multiple input masses. Relative abundance was calculated as the fraction of PacBio HiFi reads aligning to each reference genome. Each input mass was analyzed with one sequenced workflow replicate. Similar relative abundances were observed across the tested input range. **(C**,**D)** Read-length distributions shown as kernel density estimation (KDE) of PacBio HiFi data generated from DNA (C) and RNA (D) mock communities after long-read LBV-Seq. **(E**,**F)** Breadth of genome coverage for DNA viruses (E) and RNA viruses (F), shown as the percentage of each reference genome covered by at least one mapped HiFi read. Most viral genomes exhibited broad coverage. **(G)** Fraction of each viral genome recovered on individual long reads. For some small viral genomes, individual reads recovered large fractions of the genome and, in some cases, the full genome. Together, these data show that the long-read implementation generates PacBio-compatible products and supports genome detection and partial-to-complete recovery of small viral genomes on single reads.

We then sequenced the amplified products by PacBio HiFi to evaluate genome-wide performance. Mean read length was 3,677 bp across the DNA and RNA mock communities, with some reads extending to 55 kb (Fig. 3C–D). At an average depth of 575,000 HiFi reads per sample, 87.6% to 97.8% of mapped reads were classified as non-duplicate after duplicate marking, using exact sequence matches, consistent with high library complexity rather than overrepresentation of a small number of amplified molecules.

To determine whether these long reads captured broad portions of each viral genome, we aligned HiFi reads to the corresponding reference genomes. Most viruses showed high breadth of coverage (>90%) across input biomasses (Fig. 3E–F). We generally saw even coverage along the genome and calculated Gini coefficients to quantify evenness (Fig S7, Table S1). Again, AAV was the exception. As discussed previously, its predicted relative abundance (1.3 × 10^-5^) was likely too low for reliable detection at the sequencing depth used. In DNA Mock Communities 5 and 6, only a single read aligned to AAV, supporting the interpretation that its lower recovery was driven primarily by sequencing-depth limitations.

An advantage of long-read sequencing is recovery of substantial portions of viral genomes on individual reads without reference-guided reconstruction. We therefore asked how much of each genome could be captured on a single HiFi read. For several small viral genomes, including Qβ, M13, MS2, and Φ6, many individual reads spanned more than 50% of the reference genome (Fig. 3G). Notably, for M13, individual reads captured the full genome on a single molecule, and the genome could also be circularized, providing orthogonal support for genome completeness.

Overall, LBV-Seq generated PacBio-compatible products from low-biomass viral communities while preserving relative amplification behavior across input masses. These reads recovered substantial fractions of small viral genomes, and in some cases complete genomes on individual molecules, supporting reference-independent genome recovery.

### De novo assembly enables near-complete viral genome recovery from femtogram-scale inputs

We next asked whether the amplified viral genomes could be reconstructed *de novo*. This question is important because a major advantage of whole-genome sequencing is recovery of complete or near-complete viral genomes, rather than detection of only sparse genomic regions. Because assembly performance is affected by the presence of non-viral nucleic acids, we first bioinformatically removed known host sequences from the dataset. We focused the primary assembly analysis on the Illumina data because the higher read depth and co-assembly of technical replicates produced sufficient genome coverage to compare assembly performance across input masses. In contrast, the lower PacBio read depth limited genome recovery, making the PacBio datasets unsuitable for quantitative comparisons of assembly performance. We then co-assembled short-read data by combining technical replicates within each mock community. Putative viral contigs were identified with geNomad [23], which was also used for taxonomic classification and detection of inverted terminal-repeat (ITR) and direct terminal-repeats (DTR) signatures. For many viruses, geNomad classified the assembled contigs as expected and identified features consistent with known genome structures, including ITR or DTR signatures in ΦKZ, T4, M13, and vaccinia.

We next quantified assembly performance by aligning assembled contigs to the corresponding reference genomes and measuring both the number of aligned contigs and the fraction of each reference genome recovered across them (Fig. 4A,B). In the DNA mock communities, vaccinia, T4, ΦKZ, and M13 were recovered nearly completely across all input masses, typically in one or two contigs. PBCV-1 was also recovered to high completeness, but required more contigs (4-8 contigs), consistent with greater fragmentation of this larger genome. The only DNA virus we were unable to assemble well was AAV, likely due to the low sequencing depth, low relative abundance (1.3 × 10^-5^), and potentially encapsulation heterogeneity, as discussed previously, that did not enable continuous assembly. In the RNA mock communities, Qβ, MS2, and all three Φ6 segments were recovered to near-completion across all input masses, generally in one to three contigs. MHV also showed high total genome recovery but required more contigs at lower inputs, such as in Mock 5 and 6. Lentivirus also showed higher fragmentation across all input biomasses, although the majority of the genome was still assembled.

**Figure 4:**
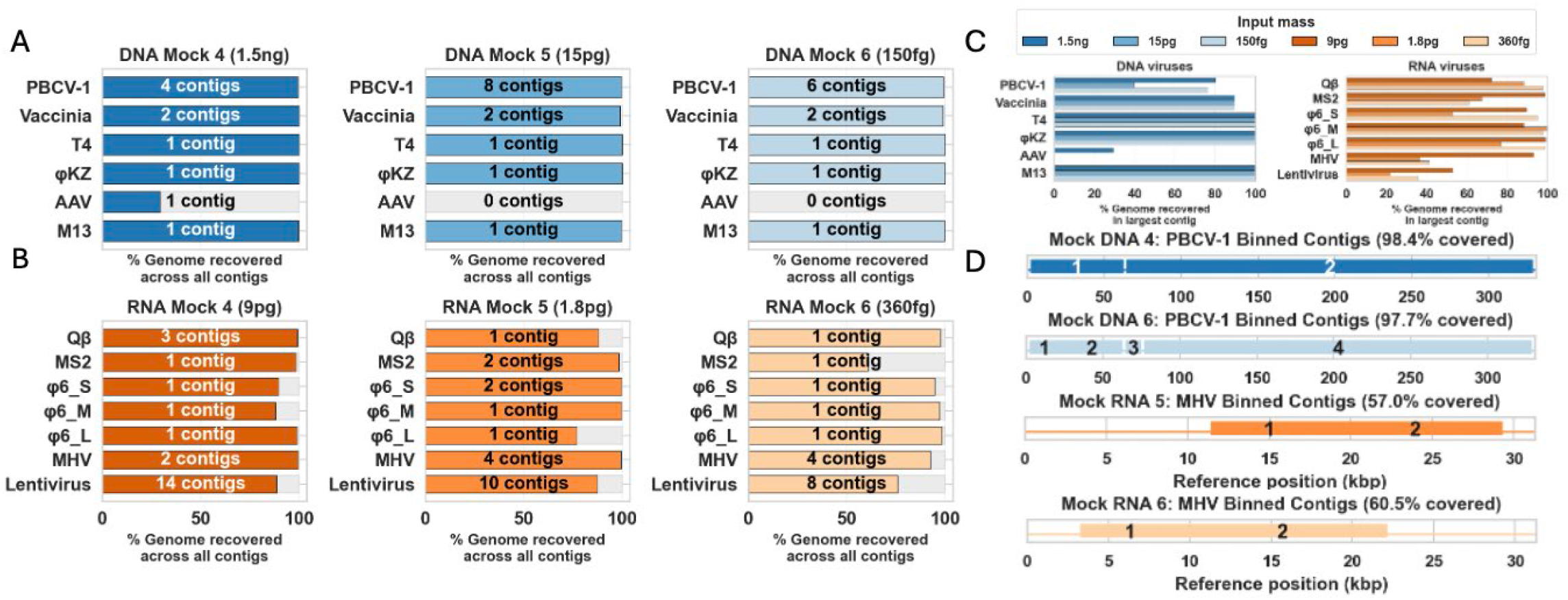
*De novo* assembly enables near-complete reconstruction of viral genomes. **(A**,**B)** Genome reconstruction from Illumina de novo assemblies were evaluated as the percentage of each viral reference genome recovered across all contigs aligning to that virus in the DNA (A) and RNA (B) mock communities. Numbers on bars indicate the number of aligned contigs contributing to that recovery. **(C)** Genome contiguity was evaluated as the percentage of each reference genome recovered in the largest *de novo* contig (defined as the contig covering the greatest number of bases in the reference genome) for each virus. **(D)** For viruses not recovered in a single contig, viral binning was used to group related contigs. Representative union plots are shown for PBCV-1 from DNA mocks 4 and 6 and MHV from RNA mocks 5 and 6. Numbers within boxes indicate individual contigs and the value in each title indicates the percentage of the reference genome recovered across the binned contigs.

To test compatibility of our workflow to reference-free viral identification, we also quantified the fraction of each genome recovered on a single contig (Fig. 4C). In the DNA mock community, PBCV-1, vaccinia, T4, ΦKZ, and M13 each reached greater than 80% single-contig completeness in at least one assembly condition. For vaccinia, the largest short-read contig spanned 89.7% of the genome but did not recover both genome termini. Additional short-read contigs contained terminal-repeat-associated sequence, yet these mapped ambiguously to both ends of the genome, consistent with incomplete resolution within the highly repetitive ITR regions rather than true end-to-end assembly [24]. To test if long-read sequencing could address these limitations, we analyzed paired PacBio sequencing and found long reads extending from each physical genome terminus through the ITRs and into unique core sequence, providing molecule-level linkage across the repeat/core junctions that was not achieved by short-read assembly alone (Fig. S8). PBCV-1 was also more difficult to recover on a single contig, highlighting potential greater challenges of assembling a large viral genome. AAV did not meet the 80% threshold of single-contig completeness, as assembly was largely unsuccessful for this virus, most likely due to the low relative abundance (Fig 4A, C).

In the RNA mock community, MHV, MS2, Qβ, and Φ6 (all segments) each reached greater than 80% completeness in at least one assembly condition (Fig. 4C). Lentivirus was a consistent exception, with less than half of the genome recovered on a single contig. Although assembly was less successful, recovery across all contigs was between 76-87% across input biomasses, indicating that much of the genome was at least present but fragmented across multiple assembled contigs (Fig. 4B,C).

Because single-contig assembly remained incomplete but total contig coverage was high for some viruses, we next asked whether viral binning could improve genome recovery while remaining reference-independent. For these cases, viral binning software, vRhyme [25], was able to group two to four related contigs and increased the fraction of the genome recovered. For example, PBCV-1 improved from 76-80% to 97-98% completeness in DNA Mock 4 and 6, and MHV improved from 36-41% to 57-60% in RNA Mock 5 and 6 (Fig. 4D). In contrast, binning provided little improvement for lentivirus, likely because current viral binning approaches, including vRhyme, rely on sequence-similarity features that are less effective for viral constructs that contain non-endogenous sequences.

Overall, these results show that *de novo* assembly enabled recovery of near-complete viral genomes across a wide range of input masses, including femtogram-scale inputs. When complete genomes were not recovered on a single contig, viral binning often extended genome recovery.

### Application of LBV-Seq to viral enriched eluate from human intestinal biopsies enables full genome reconstruction of prokaryotic and eukaryotic viral genomes at low viral masses (<0.1fg viral DNA)

To test whether the LBV-Seq workflow could recover viral genomes from clinical samples containing low viral mass, we applied both the short-read and long-read workflows to virus-enriched eluate from human duodenal biopsies. The viral enrichment protocol is described in a separate manuscript in preparation[26]. This protocol selectively depletes abundant human nucleic acids from biopsy-derived material and produces an enriched, low-biomass viral nucleic acid eluate for downstream sequencing. For this proof-of-principle analysis, we focused on DNA viruses from two participants, P1 and P2. We first estimated the total input DNA mass of these samples to determine whether they fell within the range benchmarked in the low-biomass DNA mock communities 4-6. From the DNA mock communities, time to amplification was linearly related to the log of input DNA mass (R^2^ = 0.99; Fig. 5A). Using this relationship as a standard curve (see Methods Viral mass estimations for more details), we estimated that the virus-enriched eluate yielded by the biopsy from participant P1 contained approximately 8.6 pg of total input DNA (95% prediction interval, 3.4–22 pg) and the biopsy from participant P2 contained approximately 340 fg of total input DNA (95% prediction interval, 0.13–0.91 pg), based on amplification times of 5.31 hr and 7.34 hr, respectively (Fig. 5A).

**Figure 5:**
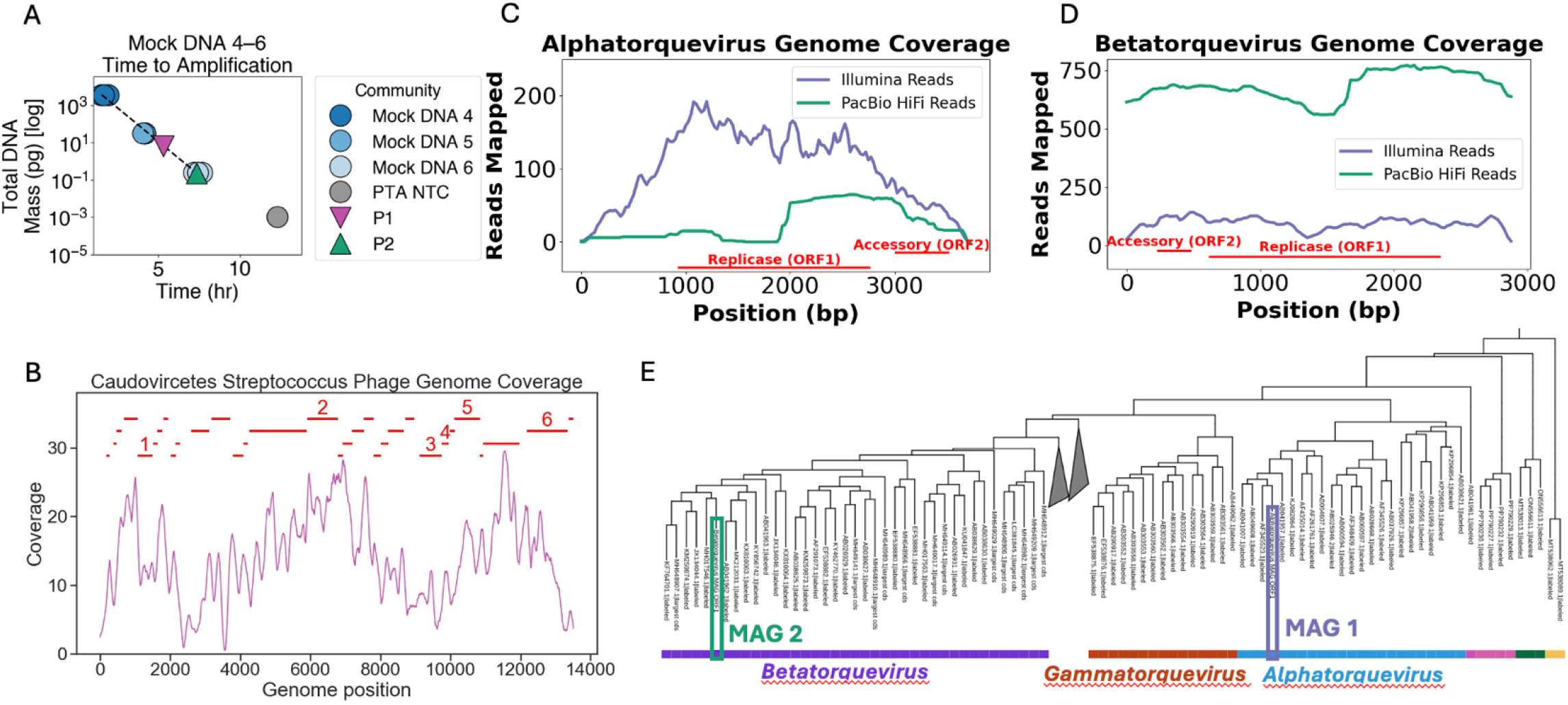
*De novo* whole genome reconstruction of prokaryotic and eukaryotic viruses from a virus-enriched eluate from a human intestinal biopsy containing low viral mass. **(A)** Total input DNA mass was estimated from time-to-amplification using DNA mock communities as a standard curve y = −1.44 log (x) + 6.65; *R*^2^ = 0.9925 (see Methods “Viral mass estimations” for more details). **(B)** Coverage of a reconstructed *Streptococcus* phage genome from Illumina short-read sequencing of a virus-enriched eluate derived from a human intestinal biopsy from participant P1 processed with LBV-Seq. Predicted open reading frames (ORFs) are shown in red, and ORFs with predicted functions are numbered. ORF1-6 have the following predicted functions: transcriptional activator, DNA polymerase/primase, transcriptional regulator, transcriptional regulator, transcriptional repressor, and integrase. **(C**,**D)** Coverage of reconstructed Alphatorquevirus and Betatorquevirus genomes in paired short-read and long-read sequencing data from the virus-enriched eluate from a human intestinal biopsy from participant P2 processed with LBV-Seq. Predicted ORFs are shown below each genome in red. **(E)** Phylogenetic placement of the reconstructed genomes using ORF1 amino acid sequences and ICTV exemplar anelloviruses, supported classification within the Alphatorquevirus and Betatorquevirus genera. Together, these data support recovery of both prokaryotic and eukaryotic viral genomes, including co-occurring anelloviruses, from virus-enriched eluates derived from human intestinal tissue processed with LBV-Seq.

We next asked whether viral genomes could be reconstructed de novo from these two low-biomass samples. In P1, we identified numerous phage contigs associated with bacterial hosts in the human oral cavity, including *Streptococcus, Veillonella, Haemophilus*, and *Neisseria* (235 phage contigs with host annotations after dereplication at <95% identity; SI Table S2). We focused on one *Streptococcus* phage genome that was classified as Caudoviricetes by geNomad and annotated as 100% complete by CheckV. This genome showed high similarity to *Streptococcus* satellite phage Javan434 (92% identity across 70% of the reference; GenBank MK448495.1), which was originally identified from the human oral cavity. This observation is consistent with prior work suggesting that the proximal small-intestine microbiome shares features with the oral microbiome [27, 28]. We further annotated open reading frames in this genome, including genes encoding integrase, transcriptional regulators and repressors, and DNA primase. Based on time-to-amplification, we estimated that this phage was present at approximately 5.3 genome copies per µL of elution (95% prediction interval, 2.1–14 copies per µL).

In P2, short-read assembly recovered two Anellovirus contigs, one classified as an Alphatorquevirus and the other as a Betatorquevirus, viruses that are considered commensal to the human virome. Long-read sequencing provided orthogonal support for both genomes. From the PacBio HiFi sequencing data, we recovered a circularized read corresponding to the same Betatorquevirus and also observed mapped long reads supporting the Alphatorquevirus genome (Fig. 5C–D), indicating the presence of two distinct anellovirus genomes, classified as Alphatorquevirus and Betatorquevirus, co-occurred in the same biopsy-derived eluate. Notably, the complete Betatorquevirus genome was also captured on a single long read, providing strong support that this genome represents a biological sequence rather than a bioinformatic artifact. Based on the time-to-amplification and percentage of sequences aligned to each viral genome constructed, we estimated abundances of 2.5 – 18 genome copies per µL (about 0.0095 fg/µL – 0.067 fg/µL) for Alphatorquevirus and 2.3 - 16 genome copies per µL (about 0.0068 fg/µL – 0.048 fg/µL) for Betatorquevirus.

The reconstructed Alphatorquevirus genome showed high similarity to known sequences in NCBI (96.7% identity across 96% of the genome; GenBank MN765522.1, BLASTn). In contrast, the reconstructed Betatorquevirus genome showed much more limited nucleotide-level similarity, with the closest match reaching only 83.3% identity across 20% of the genome (GenBank OK574447.1, BLASTn). These results suggest that the workflow can recover both viruses closely related to known references and more divergent viruses from ultra-low-input samples. To further validate these Anellovirus genomes, we predicted open reading frames and used ORF1 amino acid sequences for phylogenetic classification. In phylogenetic analysis against ICTV exemplar Anelloviruses [29], the reconstructed genomes clustered with members of the Alphatorquevirus and Betatorquevirus genera, respectively (Fig. 5E).

Finally, to assess whether these viral genomes could have arisen from background contamination, we examined the processing blanks. Alignment to the identified Anellovirus and *Streptococcus* phage genomes was minimal: seven of eight blanks showed no Anellovirus-aligned reads, one blank contained a single aligned Illumina read, and similarly little signal was observed for the *Streptococcus* phage genomes (Fig. S9).

Together, these results establish the feasibility of LBV-Seq for recovering biologically meaningful prokaryotic and eukaryotic viral genomes from human intestinal biopsies under the conditions tested, including genomes present at approximately 10 copies per µL and estimated viral masses below 0.1 fg per µL.

## Discussion

A central challenge in human virome research is to achieve recovery of viral genomes in samples that yield minimal viral nucleic acids. This challenge arises both in low-biomass and host-rich samples, including clinically relevant samples such as human tissues. Here, we developed an ultra-low-biomass virome workflow, LBV-Seq, and demonstrate that it supports *de novo*, genome-resolved analysis from samples with low viral nucleic acid mass. In both mock viral communities and virus-enriched human intestinal biopsies, this workflow supports recovery of viral genomes from nucleic acid input masses spanning from sub-femtogram to nanogram range and preserved community structure across viral input masses.

The short-read implementation of LBV-Seq provides the depth and reproducibility needed for supporting high-quality genome-resolved analysis. Long-read LBV-Seq complements Illumina sequencing by providing orthogonal evidence for genome structure, particularly in low-complexity regions such as inverted terminal repeats, and by improving completeness for small viral genomes that can be captured on single reads.

Prior methods that rely primarily on mapping or targeted capture are limited to detecting known viruses with published sequences even though many human-associated viruses are still poorly represented in current databases. Thus, the LBV-Seq workflow may provide a powerful tool for viral discovery, strain-resolved analysis, and interpretation of genome organization and gene content.

The results obtained from the virus-enriched eluate from a human-biopsy are particularly demonstrative because they validate the intended use-case that motivated this study: clinically relevant samples that are often considered too low in viral biomass for genome-resolved analysis. From virus-enriched duodenal biopsies, we recovered both bacteriophage and eukaryotic viral genomes, including co-occuring Anelloviruses at very low estimated abundance (< 0.1fg per µL). One reconstructed genome showed close similarity to known references, whereas another was substantially more divergent at the nucleotide level, illustrating that the workflow can recover both expected and less well represented viral genomes. More broadly, these results suggest that samples previously considered suitable only for detection-based virome studies may now support discovery-oriented, genome-resolved analysis.

Although the human virome was a motivating application here, the applications of the LBV-Seq workflow are much broader. In principle, the same constraints addressed in this study also arise in many low-biomass settings, including environmental, engineered, agricultural, and other host-associated samples in which standard workflows fail primarily because too little viral material is available for genome-resolved analysis.

We highlight several practical considerations for the LBV-Seq workflow. First, performance of the workflow depends on genome context. Smaller viral genomes benefit most from the long-read workflow, where long-read sequencing can completely bypass assembly requirements. Second, although the mock communities were designed to be diverse, mock communities cannot capture the full complexity of endogenous viromes, and broader validation across more compositionally complex samples is needed. Third, we did not perform a head-to-head benchmark of all whole-genome amplification chemistries, as earlier methods, particularly MDA-based approaches, are already well known to introduce strong biases in some viral settings, including preferential amplification of small circular ssDNA genomes [14–17]. Instead, we asked whether PTA could recover viral genomes reproducibly from ultra-low-input material at sufficient coverage for genome-resolved virome analysis. Within that scope, the data support LBV-Seq as a useful strategy for preserving community structure, generating broad genome coverage, and enabling *de novo* viral genome recovery from samples that would otherwise be below the threshold for such analysis. We note that an accompanying paper extends these methods to low-biomass metagenomic bacterial whole genome sequencing in human tissues, including additional validation on single nucleotide variants (SNV)-level performance in the context of Crohn’s disease [30].

Overall, LBV-Seq demonstrates a practical path toward genome-resolved analysis of low-input human virome samples. LBV-Seq shifts low-input virome analysis from viral detection toward genome-resolved interpretation. This workflow enables direct assessment of complete or near-complete viral genome reconstruction, strain-level variation, viral gene content, and genome architecture in samples with low viral mass. We anticipate that these capabilities will expand viral discovery and broaden the range of sample types amenable to genome-resolved human virome analyses. More broadly, these capabilities may facilitate deeper investigations of viral diversity, persistence, transmission, and host-associated viral ecology across animal and human states of disease and health.

## Acknowledgements

We thank Natasha Shelby for her contributions to writing and editing this manuscript. We also thank members of the lab, including Matt Cooper, Ojas Pradhan, Shruteek Marial, Matt Ratanapanichkich, Roey Lazarovits, Si Hyung Jin, Jenna Houle, and Veronica Chen, for their valuable feedback during manuscript preparation. We are grateful to Frederic Bushman, Andrew Marques, and Jiayi Duan for providing materials and guidance. We further thank Esther Leem, Rebecca Voorhees, Kaihang Wang, Raymond Zhang, Jolena Zhou, James Van Etten, Tim Shay, Erini Galatis, and the CLOVER facility for providing isolates for the mock communities. We thank Erini Galatis and Elaine Shen for their help in qPCR primer validation. We thank Scott Handley for his insightful comments and feedback during our manuscript preparation. We acknowledge Shari Kyman and the team at the TGen Collaborative Sequencing Center (CSC), as well as Xinmin Li, Hyun-Ju Lim, JaeMo Kim, and the team at the UCLA Technology Center for Genomics & Bioinformatics (TCGB), for their support with Illumina sequencing. We acknowledge Melanie Oakes and the team at the UCI Genomics Research & Technology Hub for their support with PacBio sequencing. This work was conducted in compliance with institutional biosafety (IBC) and human subjects (IRB) protocols, with approvals from the relevant committees, including contributions from Lauriane Quenee, Manali Doshi, Matt Cooper, and Ojas Pradhan. Figure 1 was created in BioRender. Wu-Woods, N. (2026) https://BioRender.com/i6ipawi

## Data availability statement

Sequencing data is available on NCBI SRA, BioProject PRJNA1457118. All non-sequencing data are available at CaltechDATA [link TBD upon journal publication].

## Author contributions

Detailed author contribution statement can be found in the SI. All authors edited and approved the final manuscript.

## Methods

### Low-biomass practices

All laboratory work was performed in a Class II biosafety cabinet (BSC). Appropriate PPE, including a lab coat, protective sleeve covers, safety glasses, and a double layer of gloves, was worn at all times. Before use, the BSC was exposed to UV light for 20 min, after which the sash was opened, and airflow was allowed to stabilize for 5 min. Interior surfaces were then sprayed with 70% ethanol and allowed to dry. High-touch surfaces within the BSC, as well as adjacent external contact areas used for reagent and sample handling, were decontaminated with a freshly made stock of 10% bleach. Any instruments to be used (e.g., pipettes, tip boxes, vortex, centrifuge, and tube rack holders) were carefully decontaminated with the 10% bleach solution. A 20-min contact time was reached before wiping surfaces and instruments to dry with paper towels. Residual bleach was then removed with distilled water, and, after drying, all surfaces and equipment were given a final spray with 70% ethanol before work commenced.

### Viral culturing & propagation

For φKZ, T4, and Qβ, phage were propagated using modified versions of previously described protocols [31, 32]. In short, each host organism was grown in LB broth overnight at 37 °C (*E. coli* MG1655 for T4, *E. coli* K12 strain hfr F-pilus + for Qβ, and *P. aeurginonsa* PA01 for φKZ). Each host organism was then back diluted to OD=0.1 before a 100 μL of high titer phage lysate was added to the cultures. Cultures were then incubated at 37 °C with agitation for ∼5 h or until lysate clears. Following this, each culture was centrifuged at 4,000 × g for 2 min. The supernatant was then filter-sterilized using a 0.4 μm filter to yield a bacterial cell-free phage lysate. Next, 0.1 volumes of chloroform were added to the supernatant, vortexed, and incubated at room temp for 10 min. This solution was then centrifuged at 4,000 × g for 5 min before the supernatant was transferred into a new tube and stored at 4 °C.

Isolate viral stocks for AAV, Phi6, vaccinia, M13, MHV, and MS2 were obtained from the Fredric Bushman group at University of Pennsylvania. Each viral stock was prepared as described previously [33]. PBCV-1 stocks were obtained from the James Van Etten group at University of Nebraska. Each viral stock was prepared as described previously [34]. Lentivirus stocks were obtained from Dr. Rebecca Vorhees group at Caltech. Each viral stock was prepared as described previously [35].

### Viral stock extractions

Viral cultures were individually processed with either filtration based VLPs or viralMEM. The choice between the VLP method was based on whether the sample readily passed through a 0.22-µm filter; samples that induced filter blockage were processed with viralMEM. For filtration based VLP enrichment, samples were vortexed briefly and passed through a 0.22µm syringe filter (Ref #SLGV004SL Millipore). The filtrate was then nuclease treated using 2 µL benzonase (EMD Millipore catalog no. 71205) and 10 µl of buffer (100 mM Tris + 40 mM MgCl2, pH 8.0 and 0.22 µm sterile filtered). Tubes were placed on a dry block incubator for 15 min at 37 °C with shaking at 600 rpm. For viralMEM processing, a modified version of a previously published protocol was performed [36].

### Mock community creation

DNA mock viral community creation 4-6: DNA mock communities were prepared from isolated genomic DNA of six viruses: PBCV-1, AAV, T4, φKZ, M13, and vaccinia virus. Viral extractions were performed (see “Viral stock extractions”). The volume of each viral DNA stock added to the mock was determined from virus-specific qPCR Cq values. DNA Mock Community 4 was generated by combining the viral DNA inputs described above. DNA Mock Community 4 contained 3.48 ng of total DNA in 3 µL of input, corresponding to 1.16 ng/µL as measured by Qubit. To quantify the viral nucleic acid mass in DNA Mock Community 4, we used virus-specific qPCR standard curves to convert Cq values to viral genome copies per reaction, as described in “Quantification: qPCR”. We then converted viral genome copies to nucleic acid mass using each virus’s genome size and genome type, accounting for whether the genome was single-stranded or double-stranded DNA. The estimated nucleic acid masses of all viral targets were summed to prepare DNA Mock Community 4 with 1.5 ng of total viral nucleic acid input. DNA Mock Communities 5 and 6 were then prepared as sequential 100-fold dilutions of DNA Mock Community 4, yielding estimated viral DNA input masses of 15 pg and 150 fg, respectively. Mock communities were prepared at sufficient volume for triplicate viral quantification measurements and triplicate LBV-Seq reactions, with remaining material reserved for downstream applications.

RNA mock viral community creation 4-6: RNA mock communities were prepared from isolated genomic RNA of five viruses: MHV, MS2, Qβ, φ6, and lentivirus. Viral extractions were performed (see “Viral stock extractions”). The volume of each viral RNA stock added to the mock was determined from virus-specific qPCR Cq values.

To reduce host-derived background signal, original Qβ phage and MS2 phage stocks were treated with TURBO DNase (Cat #AM2238; Thermo). MHV and lentivirus stocks were subjected to mammalian and bacterial rRNA depletion from NEB (Cat #E7400S & E7850S) according to the manufacturer’s protocol using mixed bacterial and mammalian probe sets. To quantify the viral nucleic acid mass in RNA Mock Community 4, we used virus-specific qPCR standard curves to convert Cq values to viral genome copies per reaction, as described in “Quantification: qPCR”. We then converted viral genome copies to nucleic acid mass using each virus’s genome size and genome type, accounting for whether the genome was single-stranded or double-stranded RNA. We summed the estimated nucleic acid masses across all viral targets to prepare RNA Mock Community 4 with 9 pg of total viral nucleic acid input. RNA Mock Communities 5 and 6 were then prepared by serial 5-fold dilution of RNA Mock Community 4, yielding estimated viral RNA input masses of 1.8 pg and 360 fg,. Reverse transcription was performed in triplicate for each mock community using SuperScript IV (Cat # 18090200; Thermo), with one no-template control included. Following reverse-transcription, cDNA was purified using AMPure XP beads (Cat # A63881; Beckman Coulter) at a 1.8× bead-to-sample ratio and eluted in 20 µL. The purified products were used for downstream analyses, including post-RT quantification by qPCR.

DNA mock viral community creation 1-2: DNA mock communities 1 and 2 were prepared from purified viral genomic DNA. DNA Mock 1 contained six viruses, PBCV-1, AAV, T4, φKZ, M13, and vaccinia virus. DNA Mock 2 contained only double-stranded DNA viruses, PBCV-1, T4, φKZ, and vaccinia virus. The volume of each viral DNA stock added to each mock community was determined from virus-specific qPCR Cq values. In DNA Mock 1, target relative abundances by genome copy were 36% φKZ, 35% vaccinia virus, 20% T4, 5% PBCV-1, 3% M13 phage, and 1% AAV. Viruses shared between DNA Mock 1 and DNA Mock 2 were added at the same target genome-copy abundances. After mock communities were generated, LBV-seq short-read workflow was performed in triplicate from each mock community.

### Human sample processing

All activities related to the enrollment of the two participants, collection of samples, and sample analyses were approved by the University of Chicago and Mayo Clinic IRB committees and performed under IRB protocols 22-1138-AM023 and 22-007133 respectively. Deidentified samples were received at Caltech and analyzed under Caltech IRB protocol IR22-1271, a reliance on the UC protocol, and Caltech IRB #19-0915, an exempt protocol which allows for secondary use of biospecimens originally obtained from other studies.

For the purposes of this study, we selected two participants whose biopsies contained detectable viral loads. The two participants were celiac disease patients and were consented and recruited at the Mayo Clinic or the University of Chicago to participate in a separate study examining the responses of newly diagnosed celiac disease patients to a gluten-free diet. The two participants were screened for diagnosis and eligibility criteria for enrollment in the study. Exclusion criteria included: participants with chronic infectious diseases such as human immunodeficiency virus or hepatitis C (HCV); active, untreated *Clostridium difficile* infection; active infection with severe acute respiratory syndrome coronavirus 2 (SARS-CoV-2); intravenous or illicit drug use such as cocaine, heroin, nonprescription methamphetamines; active use of blood thinners; severe comorbid diseases; participants on active cancer treatment and participants who were pregnant. Approaching prospective participants was at the discretion of their treating physician and was not done in cases that would put participants at any increased risk, regardless of reason. Participants underwent upper endoscopy with either moderate or deep sedation. Duodenal biopsies were collected during an endoscopy and flash frozen in liquid nitrogen in the clinic. The duodenal biopsies were stored at -80°C until shipment to Caltech, which was done using dry ice. Upon receipt by Caltech, they were processed with viralMEM, a modified version of the protocol here was performed [36]. The viral fraction was extracted following the protocol outlined in “Viral stock extractions”.

### LBV-Seq amplification

LBV-Seq for short-read sequencing: 3 µLs of mock community mixtures of gDNA were transferred into 96-well plate in triplicate. Modifications of ResolveDNA Whole Genome Amplification Kit v1 (Bioskryb, PN: 100500) protocol were performed due to not needing to lyse cells and to limit DNA modifications due to harsh alkaline lysis chemistry [21]. 3 µL of gDNA were added to MS mixtures (omiting cell lysis buffer), incubated at 10 °C for 10 min. SB4 mastermix was modified to include 0.1µL of 20X EvaGreen dye (BioRad Laboratories) to facilitate real-time tracking and amplification time was extended to 14 hours. All reagent and mastermixes were added by an EpMotion5075 protocol (Eppendorf), as well as on-board incubations described by the manufacturer’s protocol. Plates were loaded into a Bio-Rad CFX96 and incubated for 168 cycles of 5 min per cycle, with a plate read after each cycle, followed by a 12 °C hold. The amplified DNA was subsequently cleaned using 2X AMPure XP beads (Cat # A63881; Beckman Coulter) and utilized in “Quantification” and “Illumina Library Prep & Sequencing”.

LBV-Seq for long-read sequencing: Extracted DNA was transferred into 96-well plates using 3 µL per sample. The plate was shipped on dry ice to BioSkryb for PTA long-read amplification, termed the Services Custom PacBio Long Read Amplification (BioSkryb Genomics 101157). In brief, the ResolveDNA Whole Genome Amplification Kit was modified to increase amplified DNA size. The received DNA was cleaned using 2X AMPure XP beads and QC’d with Tapestation and Qubit double-stranded DNA (dsDNA) High Sensitivity assay (Thermo Cat #Q32851). Subsequent DNA was used as described below in the sections “Quantification” and “PacBio library prep & sequencing”.

### Quantification

qPCR: Quantification for each viral target part of the Mock Communities was quantified using quantitative PCR (qPCR) on the Bio-Rad CFX96 Thermocycler. Primers for each viral target were designed and validated.

Annealing temperature was optimized using a temperature gradient. Optimized annealing temperatures for each target are reported in (SI Table S3). For each mock viral community, we quantified the concentration of each viral target using target-specific qPCR thresholded at 339 relative fluorescent units (RFU). For each target, we generated a standard curve using purified amplicons serially diluted 10-fold from 1 × 10^9^ to 1 × 10^−1^ copies per reaction, with each dilution series performed in triplicate. We fit a linear regression to the standard curve over the reliable quantitative range and used this regression to convert sample Cq values to target copies per reaction (Fig S10). We then converted target copies per reaction to estimated genome copies per microliter of input sample, assuming one qPCR target copy per viral genome. To estimate the number of genome bases contributed by each viral target, we multiplied the genome copies per microliter by the length of that viral genome. We calculated the expected relative abundance of each viral target as the fraction of total genome bases contributed by that target across all targets in the mock community. Finally, we estimated the nucleic acid mass contributed by each target by multiplying the number of genome bases by the mass of one base pair, 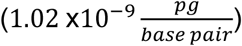) [37].

#### Qubit

Mock community DNA 4 was quantified with the Qubit double-stranded DNA (dsDNA) High Sensitivity (HS) Assay Kit (Thermo Cat #Q32854) on a Qubit 3 Fluorometer. Working solution was prepared by diluting the dsDNA HS reagent 1:200 in the supplied buffer, and 2 µL of sample was added to 198 µL of working solution. Samples were briefly vortexed and incubated for 2 min at room temperature. Sample concentration was measured following calibration with supplied standards according to the manufacturer’s instructions and yielded 1.16 ng/µL in DNA Mock Community 4.

Human-derived viral mass estimations: DNA Mock Community 4-6 was used to generate a standard curve because the total DNA input mass of this community was known and quantifiable. DNA Mock Communities 5 and 6 were treated as 100-fold and 10,000-fold dilutions of DNA Mock Community 4, respectively. For each mock community, amplification time was measured for three technical replicates by qPCR on a Bio-Rad CFX96. The fluorescence threshold was set at 202 relative fluorescence units (RFU). Cq values were converted to amplification time by multiplying each Cq value by 5 min, corresponding to the duration of each cycle. A standard curve was generated by plotting amplification time against total DNA input mass. Linear regression yielded the equation y = −1.44 log(x) + 6.65, with R^2^ = 0.9925, where x represents total DNA input mass in picograms and y represents amplification time. For virus-enriched eluates from human clinical samples, total DNA input mass could not be directly quantified as it is below the limit of traditional total DNA quantification measurements, such as Qubit. Amplification time was therefore measured using the same qPCR protocol, and total DNA input mass was estimated from the standard-curve equation. The DNA mass represented by each assembled viral genome was estimated from the fraction of sequencing reads assigned to that genome. For each sample, the read fraction was calculated as the number of reads mapped to a given assembled viral genome divided by the total number of sequencing reads from that sample. The read fraction was then multiplied by the estimated total DNA input mass to estimate the DNA mass represented by that assembled viral genome. The prediction interval for the total mass in each sample was calculated in the following way. First, an ordinary least-squares linear regression standard curve relating amplification time to the log-transformed total mass data. For each sample, the predicted log mass was calculated from the fitted regression line, and a 95% prediction interval was computed as 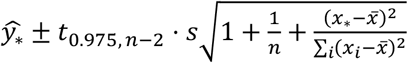 where 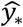 is the predicted log mass based on the amplification time *x*_*_, *s* is the residual standard error of the regression, and *n* is the number of standard-curve measurements. Predicted values and the interval bounds were then transformed from log space to obtain estimates and corresponding 95% prediction intervals on the original mass scale.

### Illumina library prep & sequencing

Amplified DNA was prepared for sequencing using the Illumina DNA Prep kit (Cat #20018704), with a maximum input of 500 ng of cleaned DNA per sample. DNA input was quantified after clean-up using the Qubit dsDNA High Sensitivity assay (Thermo Fisher Scientific, Cat #Q32854). Final libraries were evaluated for fragment size and overall quality using a High Sensitivity D1000 TapeStation chip (Agilent, Cat #5067-5585 and #5067-5584). Library concentrations were determined by qPCR on a CFX96 instrument (Bio-Rad, Cat #1855196) using primers targeting the Illumina adapter sequences (forward: AAT GAT ACG GCG ACC ACC GA; reverse: CAA GCA GAA GAC GGC ATA CGA). Prior to amplification, libraries were diluted 1:10,000 in nuclease-free water to ensure concentrations fell within the range of the KAPA Library Quantification standards (Roche, Cat #07960387001). Each 10 uL qPCR reaction contained 1x SsoFast EvaGreen Supermix (Bio-Rad, Cat #1725201), 250 nM forward primer, and 250 nM reverse primer.

Thermocycling conditions were 95 °C for 5 min, followed by 40 cycles of 95 °C for 30 s and 60 °C for 45 s. Pooled libraries were quantified again using the Qubit dsDNA High Sensitivity assay and assessed on a High Sensitivity D1000 TapeStation chip before submission for sequencing. Sequencing was performed at the UCLA Technology Center for Genomics & Bioinformatics (TCGB) and the TGen Collaborative Sequencing Center (CSC) on the Illumina NovaSeq X platform using 2 × 150 bp paired-end reads. Samples were demultiplexed on the NovaSeq X instrument, and raw FASTQ files for read 1 and read 2 were generated for each sample.

### PacBio Library prep & sequencing

Up to 18 µL of amplified DNA was utilized for PacBio library preparation (inputs ranged from 536 – 1,534ng). Samples were prepared using the SMRTbell® prep kit 3.0 (PacBio PN: 102-182-700) without any size selection or SRE (short read eliminator) steps. Processing began at step 4: Repair and A-tailing. Samples were processed individually until after nuclease treatment and clean-up.

Samples were subsequently QC’d with a Genomic TapeStation Chip and Qubit’s dsDNA High Sensitivity assay before pooling at equivalent masses. The pool was shipped to UCI Genomics Research and Technology Hub (GRT Hub) for processing with ABC (Annealing, binding, and clean-up) and sequencing of the pool on one SMRT Cell on the PacBio Revio system for 24 hours.

### Bioinformatic processing

Short-read processing: After receiving de-multiplexed reads from the NovaSeqX, raw fastq files were provided for r1 and r2. Adapter trimming and quality control was performed with cutadapt (v4.4) [38] and QC’d with multiQC (v1.30) [39]. For the mock viral communities 4-6, classification was performed with bowtie2 (v2.5.5) [40] alignment against a reference containing the known viruses present in the sample using default alignment parameters. Relative abundance and breadth of coverage was assessed with samtools coverage (v1.23.1) [41] using the “numreads” and “covbases” columns respectively. Assembly was performed using megahit (v1.2.9) [42] and viral sequences were identified with genomad (v 1.11.2) [23] and virsorter2 (version 2.2.4)[43]. Viral binning was performed with vRhyme (v1.1.0) [25]. Assembly completion was assessed by aligning contigs back against the viral reference sequences with minimap2 (v2.30) [44]. For characterization of host background, used in SI Fig S1, alignment in mock DNA viral communities was performed with bowtie2 (v2.5.5) [40] using a reference genome containing all input viruses and their known hosts with default alignment parameters. For mock RNA viral communities, alignment was performed with salmon (v1.10.3) [45] with default alignment parameters (including the validate Mappings flag).

For human derived sequencing (i.e. from the human intestinal biopsies), host filtering was first performed on reads through alignment to the T2T-chm13 reference genome (v2.0) [46] with bowtie2 (v2.5.5) [40] and removing any paired end read that mapped, either primary, secondary, or supplementary. For phage identification from the human samples, viral contigs were assembled using either the SPAdes genome assembler (v4.2.0) [47, 48] (in rnaviralSPAdes mode and metaviralSPAdes mode) or megahit (v1.2.9) [42]. Viral sequences were then identified with genomad (v 1.11.2) [23] and virsorter2 (version 2.2.4) [43]. The host organisms for viruses identified as phages by geNomad based on taxonomy, and with a viral score greater than 90 from either VirSorter or geNomad, were predicted using iPHoP (version 1.4.2) [49] with a minimum score threshold of 90. The completeness of viral contigs was estimated using checkV (v1.0.3)[50] and the v1.5 database with the default settings defined by the end_to_end command. Coding sequences for phage shown in Figure 5C were then annotated using pharokka (v1.9.1) [51] using the version 1.8.0 database.

Long-read processing: After receiving de-multiplexed reads from the Revio SMRT cell, raw fastq and fasta files were provided. Kernal density estimation (KDE) plots of read-length were made using the default seaborn v 0.13.2 kernel and bandwidth parameters (Fig 3C-D). For the mock viral communities 4-6, viral classification was performed with minimap2 (v2.30-r1287) [44] alignment against a reference containing the known viruses present in the sample using default alignment parameters. Relative abundance and breadth of coverage was assessed with samtools coverage (v1.23.1) [41] using the “numreads” and “covbases” columns respectively. Assembly was performed using hifiasm meta (v 0.13-r308) [52] and viral sequences were identified with genomad (v 1.11.2) [23] and virsorter2 (version 2.2.4) [43]. Assembly completion was assessed by aligning contigs back against the viral reference sequences with minimap2 (v2.30-r1287) [44].

To analyze the sequencing results from the processed human intestinal biopsies, bioinformatic host-filtering was first performed on reads through alignment to the T2T-chm13 reference genome (v2.0) [46] with minimap2 (v2.30-r1287) [44].

