## Supplementary Information for "Ultra-low biomass sequencing workflow (LBV-Seq) enables de novo metagenomic reconstruction of DNA and RNA viral genomes"

**Contents:**

Fig S1-10

Tables S1-S3

Detailed Author Contributions

Author Contact Information and ORCiDs

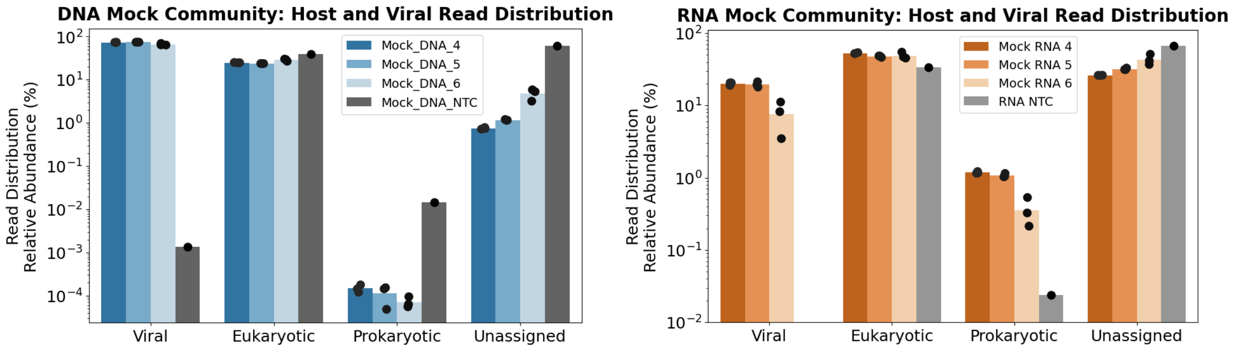


**SI Figure 1: Mock viral communities contain residual host nucleic acids due to incomplete virion preparations.** Quality-controlled reads were mapped against a reference database containing all potential reference sequences of nucleic acid present in the sample. This includes the viral and host genomes for DNA Mock communities and viral and host genomes and transcriptomes for RNA Mock communities. For DNA alignment, bowtie2 was performed and reads were summed within the domains and normalized to the total number of reads. For RNA alignment, reads against each viral host were quantified using Salmon. Significant host carryover was detected in all mock communities. DNA and RNA Mock 4 contained sufficient biomass to be quantified using Qubit, and this measurement was used as the ground-truth estimate of total mass for subsequent analyses (Fig 5A).

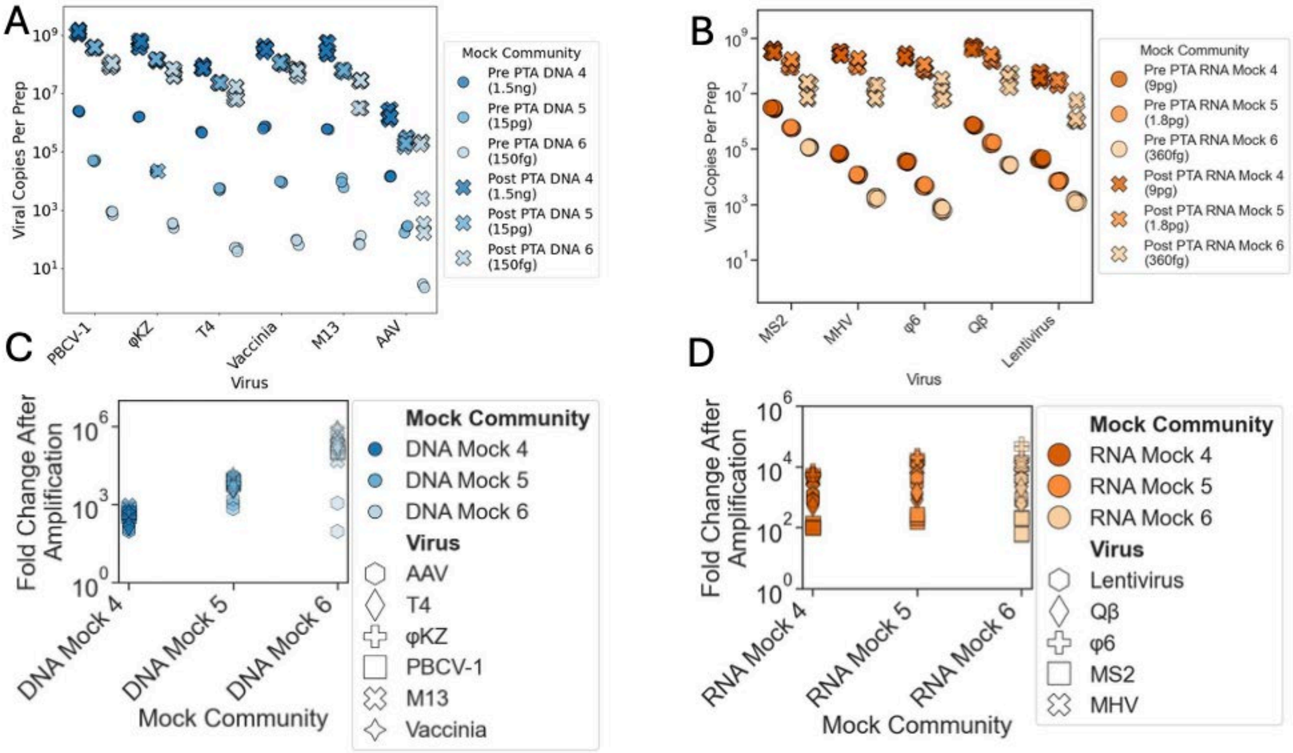


**SI Figure 2: LBV-Seq enables consistent amplification of viral genomes across biomasses. A-D)** Viral concentrations were measured by qPCR using virus-specific primers targeting single-copy genomic regions before and after amplification **(A-B).** Pre-amplification quantification is denoted by (o) and post-amplification quantification by (x). The expected viral copies into prep were calculated by multiplying qPCR estimates of copy number per µL by the input volume (3µL). The fold change in viral copies into prep was calculated by dividing the average viral copies into prep after amplification across qPCR replicates by the average viral copies into prep before amplification across qPCR replicates **(C-D)**. The fold change of each virus is denoted by a unique shape.

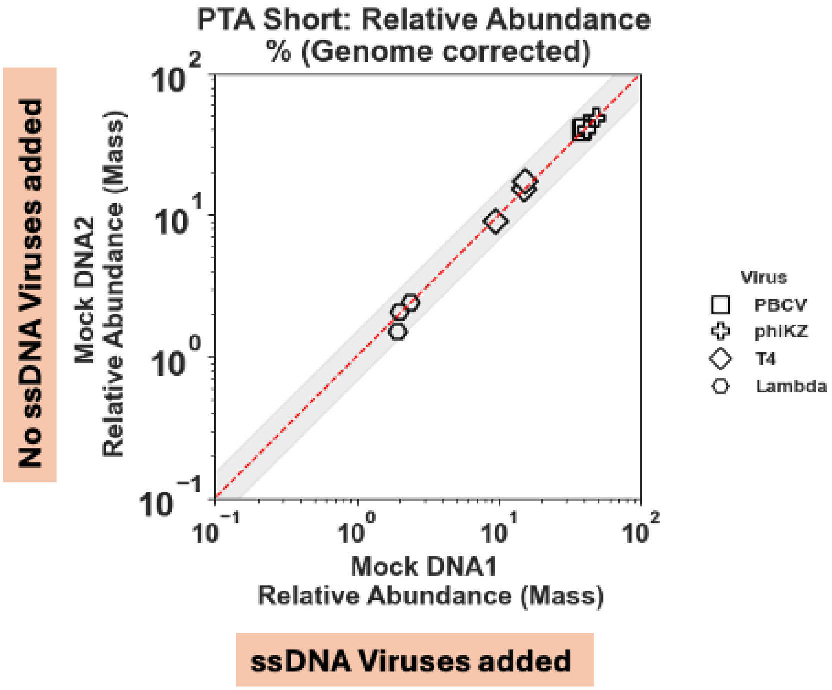


**SI Figure 3: LBV-Seq preserves dsDNA community composition in the presence of ssDNA viruses.** The short-read PTA workflow was applied to two DNA mock communities (Mock DNA 1 and Mock DNA 2) to evaluate whether ssDNA viruses introduce amplification bias. Mock DNA 1 contained a mixture of dsDNA and ssDNA viruses, including AAV and M13 (see Methods "DNA mock viral community creation 1-2"). Mock DNA 2 was identical in composition but contained only dsDNA viruses. Following sequencing, relative abundances were calculated and compared between the two communities. The dsDNA viral relative abundances were highly consistent between Mock DNA 1 and Mock DNA 2, indicating that the presence of ssDNA viruses does not substantially bias PTA-based amplification of dsDNA community composition.

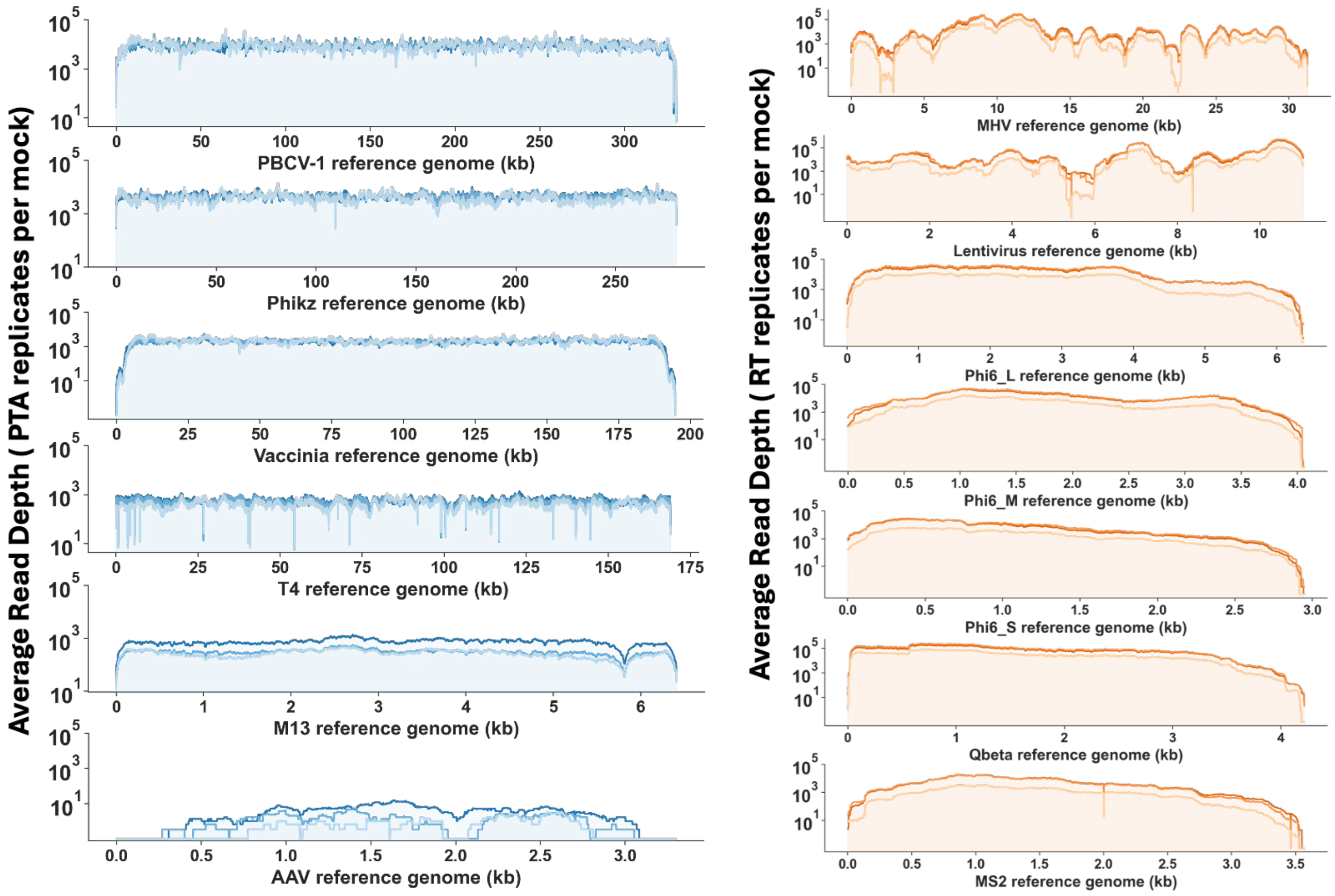


**SI Figure 4: LBV-Seq supports high breadth of coverage across diverse viral genomes over the biomass range tested.** Average read depth across reference genomes is shown with a pseudocount of 0.1X coverage for viruses in the DNA and RNA mock communities following PTA and Illumina sequencing . Coverage extended across most viral genomes, indicating that LBV-Seq supported whole-genome amplification across the tested biomass inputs. Coverage profiles were generally uniform across viruses, although coverage decreased near the ends of some genomes. These decreases occurred in viruses with long or complex inverted terminal repeats, such as Vaccinia virus and AAV, and in structured RNA genomes, such as Qβ and MS2, suggesting that terminal genome features may reduce coverage in some cases.

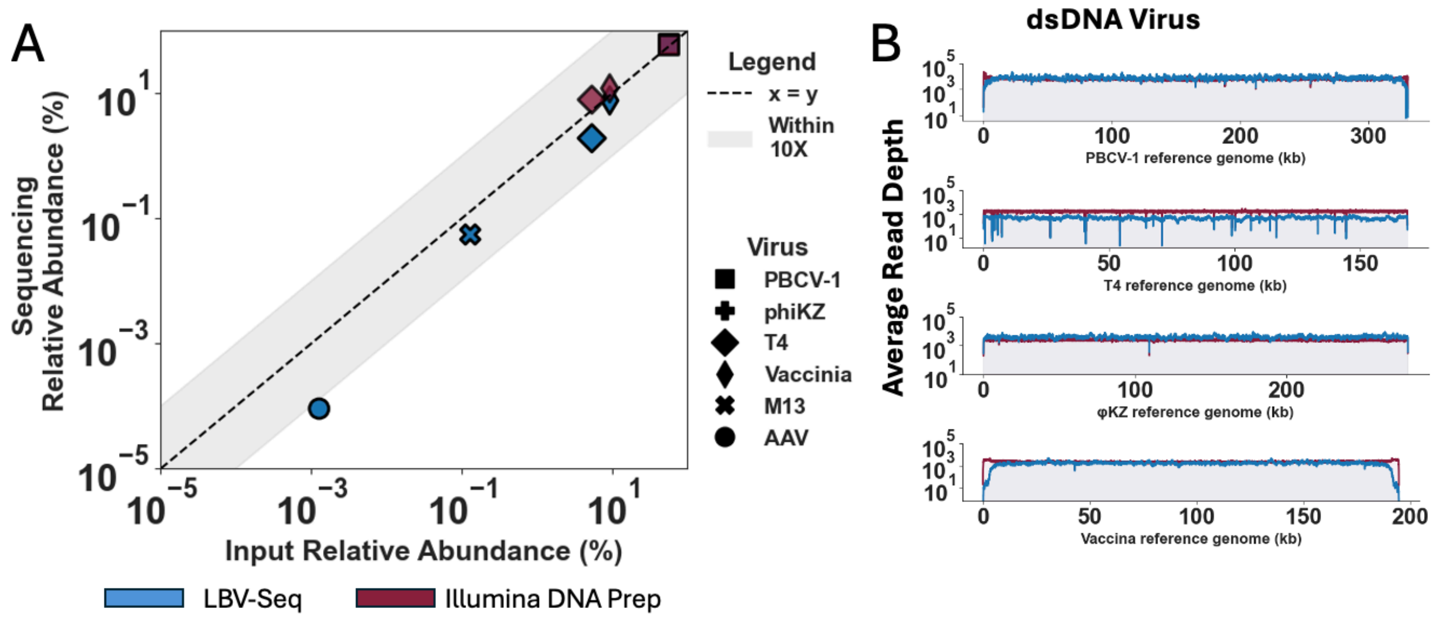


**SI Figure 5: LBV-Seq preserves dsDNA viral community relative abundance compared to Illumina DNA Prep.** DNA Mock Community 4 provided sufficient input for library preparation without whole-genome amplification, enabling direct comparison between LBV-Seq and Illumina DNA Prep. (A) After short-read Illumina sequencing, viral relative abundance was calculated as the fraction of total aligned bases assigned to each reference genome. These values were compared with the pre-amplification viral community composition measured by single-copy virus-specific qPCR assays. LBV-Seq and Illumina DNA Prep showed similar dsDNA viral community compositions relative to the qPCR-measured input community. LBV-Seq is shown in blue, and Illumina DNA Prep is shown in red. ssDNA viruses were not plotted for the Illumina DNA Prep condition because this library-preparation strategy does not efficiently capture ssDNA genomes. Shapes indicate viral targets, and gray shading indicates a 10-fold range around the expected relative abundance. (B) Coverage depth across each reference genome is shown with a pseudocount of 0.1X coverage. Genome coverage profiles were similar between LBV-Seq and Illumina DNA Prep libraries, with many peaks and troughs occurring at similar genomic positions. Some differences were observed near viral genome ends, as can be seen with the vaccinia virus.

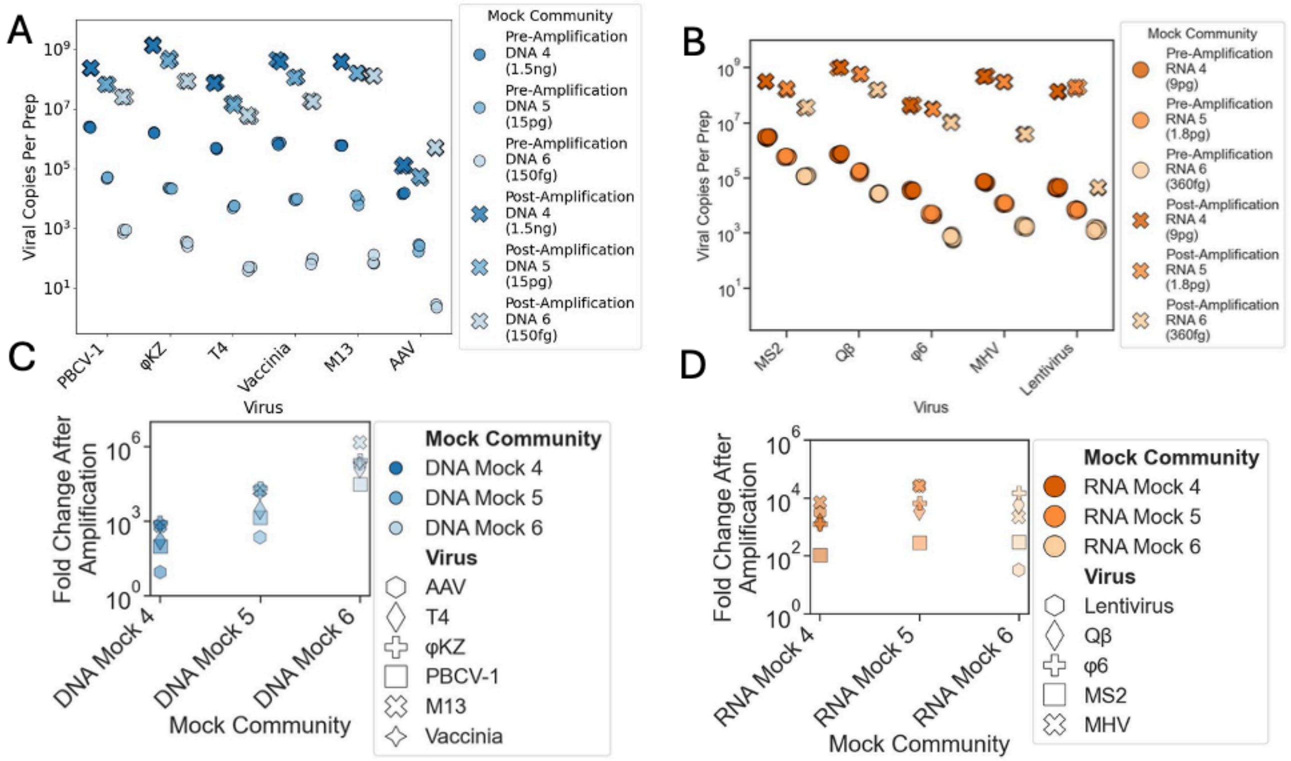


**SI Figure 6: LBV-Seq with long read-sequencing enables consistent amplification of viral genomes across biomasses. A-D)** Viral concentrations were measured by qPCR using virus-specific primers targeting single-copy genomic regions before and after amplification for long-read seqeuncing **(A-B).** Pre-amplification quantification is denoted by (o) and post-amplification quantification by (x). The expected viral copies into prep were calculated by multiplying qPCR estimates of copy number per µL by the input volume (3µL). The fold change in viral copies into prep was calculated by dividing the average viral copies into prep after amplification across qPCR replicates by the average viral copies into prep before amplification across qPCR replicates **(C-D)**. The fold change of each virus is denoted by a unique shape.

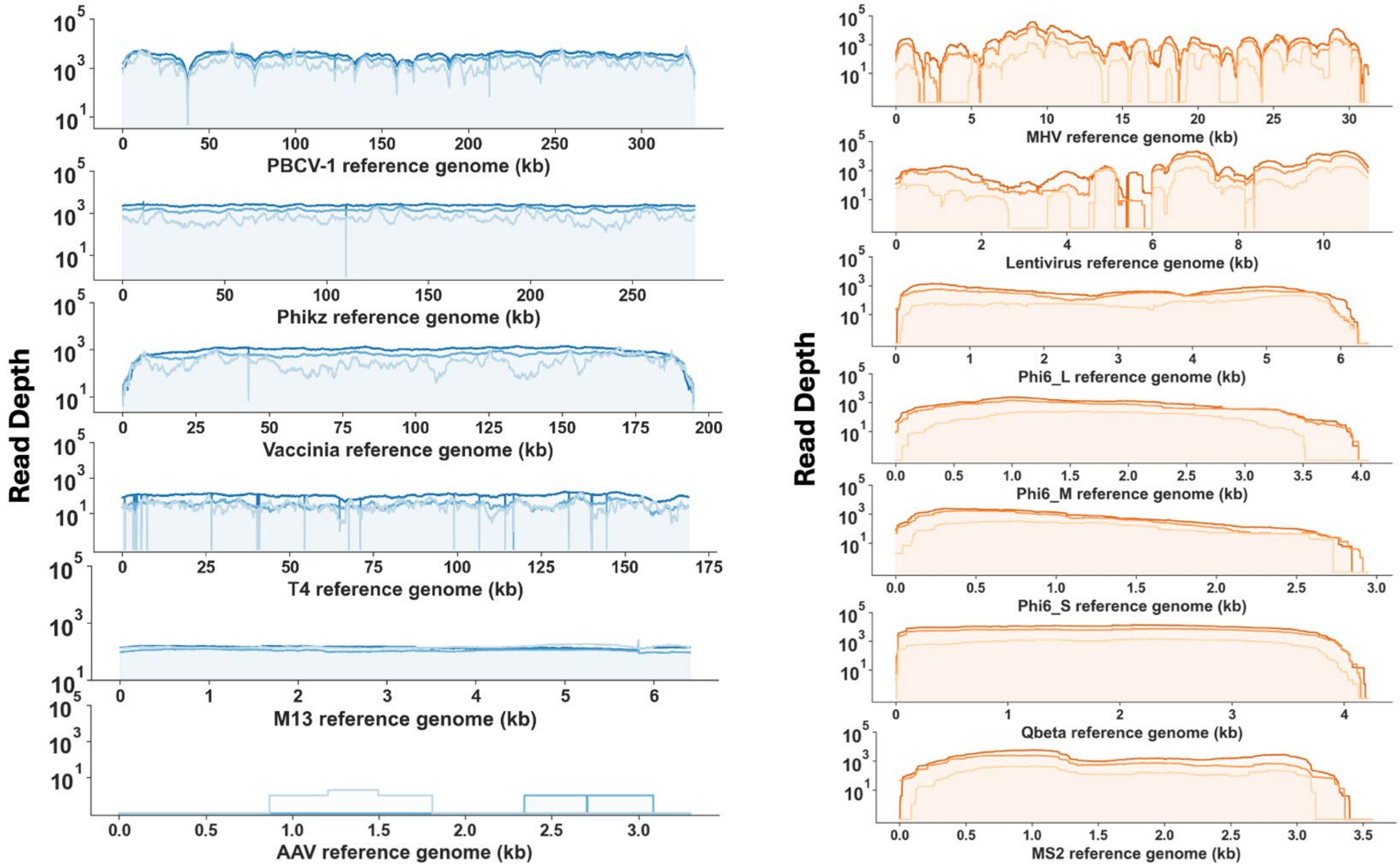


**SI Figure 7: LBV-Seq with long-read sequencing has comparable genome coverage to short read across diverse viral genomes over the biomass range tested**. Coverage depth across each reference genome is shown with a pseudocount of 0.1X coverage for viruses in the DNA and RNA mock communities after LBV-Seq for long-read and PacBio HiFi sequencing. Coverage extended across most viral genomes and input masses, indicating that the workflow generated genome-wide long-read coverage rather than sparse viral detection. As observed with short-read LBV-Seq, coverage decreased near the ends of some genomes, but reads still extended into these terminal regions.

**SI Figure 8: Long reads anchor vaccinia terminal-repeat sequence to the unique genomic core. (A)** Left terminus and **(B)** right terminus of the vaccinia genome from Mock DNA 4 community. The largest short-read contig, shown in black, spans 89.7% of the genome but breaks within the inverted terminal repeats (ITRs; shaded blue). Additional short-read contigs, shown in grey, map ambiguously to both termini. In contrast, long reads, shown in blue, extend from each physical genome end past the ITRs into unique core sequence, supporting placement of terminal-repeat sequence that is not resolved by short-read assembly alone.

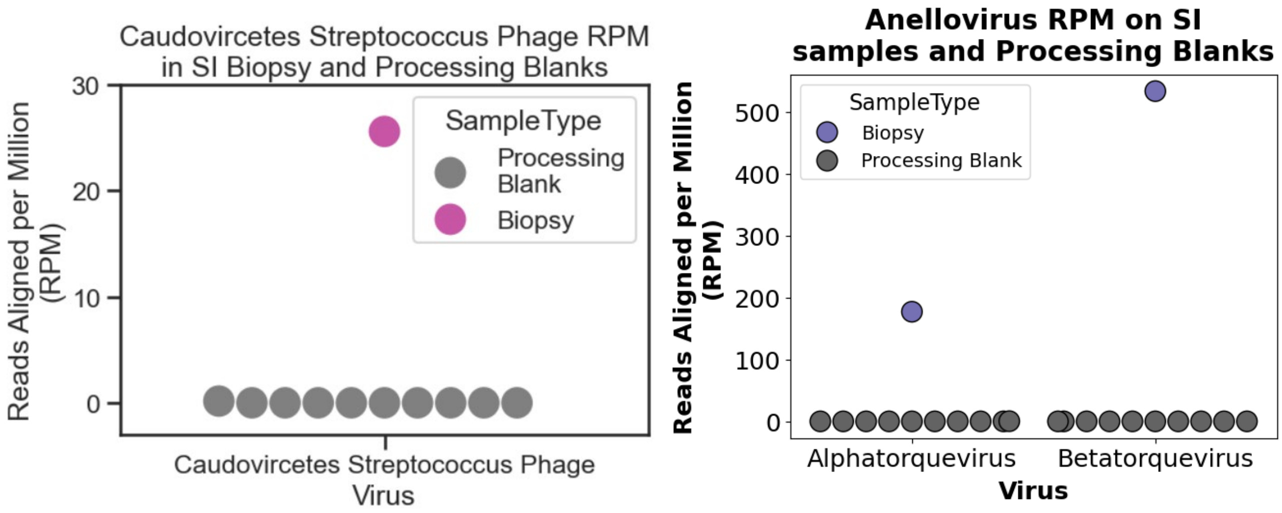


**SI Figure 9: Minimal viral signatures detected in processing blank samples.** To distinguish biological signal from background noise, the abundance of Streptococcus phage **(A)** and Anellovirus **(B)** was quantified in processing blanks (N = 8) and compared with biopsy samples. Reads from blank samples were mapped to the corresponding viral contigs, and aligned reads were normalized as reads per million (RPM). Minimal viral signal was detected across all blank samples.

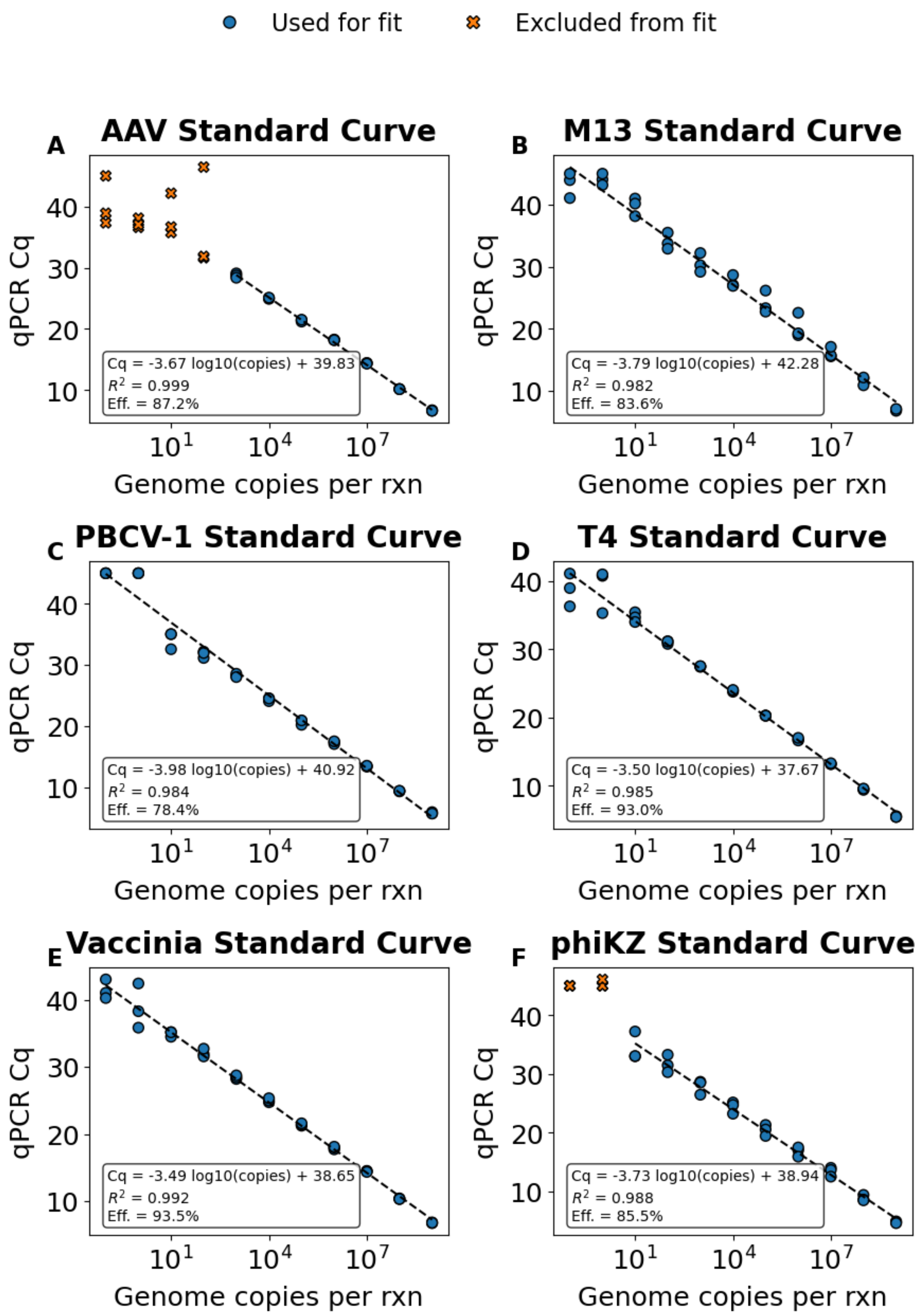
Used for fit
Excluded from fit
A
AAV Standard Curve
Cq = -3.67 log10(copies) + 39.83
$R^2$ = 0.999
Eff. = 87.2%
B
M13 Standard Curve
Cq = -3.79 log10(copies) + 42.28
$R^2$ = 0.982
Eff. = 83.6%
C
PBCV-1 Standard Curve
Cq = -3.98 log10(copies) + 40.92
$R^2$ = 0.984
Eff. = 78.4%
D
T4 Standard Curve
Cq = -3.50 log10(copies) + 37.67
$R^2$ = 0.985
Eff. = 93.0%
E
Vaccinia Standard Curve
Cq = -3.49 log10(copies) + 38.65
$R^2$ = 0.992
Eff. = 93.5%
F
phiKZ Standard Curve
Cq = -3.73 log10(copies) + 38.94
$R^2$ = 0.988
Eff. = 85.5%
qPCR Cq
Genome copies per rxn

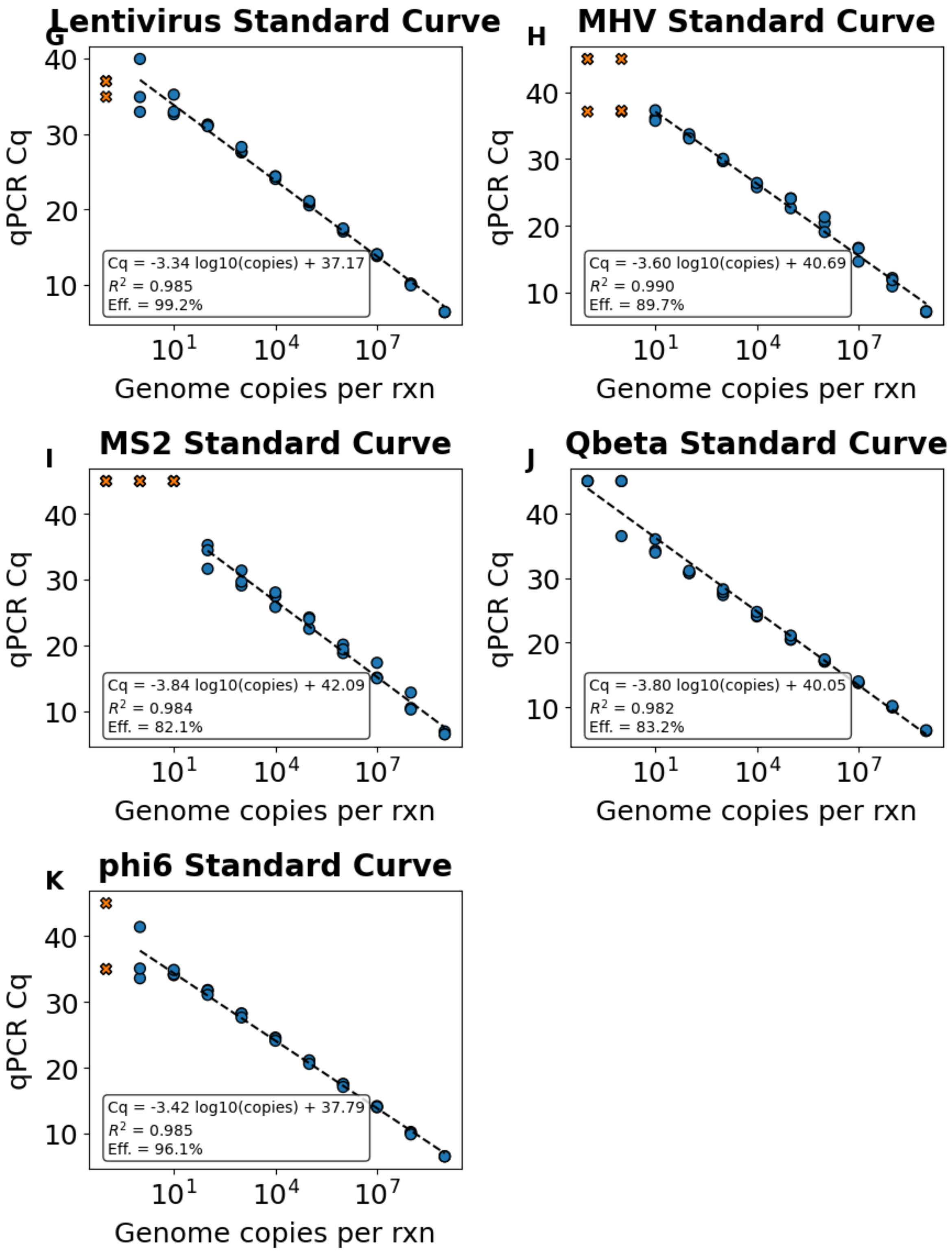


**SI Figure 10. Standard curves for each viral single-copy target qPCR assay.** Standard curves are shown for each primer set targeting DNA viruses first, followed by RNA viruses: **A**, AAV; **B**, M13; **C**, PBCV-1; **D**, T4; **E**, Vaccinia; **F**, phiKZ; **G**, Lentivirus; **H**, MHV; **I**, MS2; **J**, Qbeta; and **K**, phi6. Genome copy equivalents per reaction were plotted against qPCR Cq values. Blue circles indicate points included in the linear regression, orange X symbols indicate low-concentration points excluded from the fit, and dashed lines indicate fitted standard curves. Insets report the regression equation, coefficient of determination ($R^2$), and amplification efficiency for each assay.

**SI Table 1: Genome coverage uniformity metrics for reference-based alignments of mock viral communities sequenced using Illumina and PacBio platforms.** Coverage uniformity was evaluated using the Gini coefficient and coefficient of variation (CV) calculated from per-base read depth across each viral reference genome. Lower Gini and CV values indicate more uniform sequencing coverage, whereas higher values reflect increased coverage bias and uneven read distribution. Mean sequencing depth (mean_depth_avg) represents the average per-base coverage across the viral genome. Metrics are summarized by library construction method (LC_Method), mock community (mock_id), nucleic acid type (NA), and viral species (Virus). Standard deviation values were calculated when replicate sequencing datasets were available.

**SI Table 1 is attached as a .csv file**

**SI Table 2. Predicted bacterial hosts of potential bacteriophage contigs from participant P1.** Likely bacteriophage contigs assembled from participant P1 were assigned bacterial hosts using iPHoP. The table includes 235 non-redundant contig clusters with high-confidence genus-level host predictions, defined as iPHoP confidence scores greater than 90. Because multiple assemblers were used, closely related contigs may represent the same viral genome. For each representative contig, the table reports predicted host taxonomy, iPHoP confidence score, host-assignment support, related contigs with at least 95 percent sequence identity, and average amino acid identity to the closest RaFAH reference.

| Target | Forward Primer Sequence (5' -> 3') | Reverse Primer Sequence (5' -> 3') | Amplicon Size (bp) | Annealing Temperature (℃) |
|---|---|---|---|---|
| AAV2-GFP | GAACCGCATCGAGCTGAA | TGCTTGTCGGCCATGATATAG | 111 | 67 |
| Lentivirus-BFP | GAGAGTCACCACATACGAAGAC | CCCTCTGATCTTGACGTTGTAG | 97 | 60 |
| MS2 phage | GCTGAATGGATCAGCTCTAACT | CAGTCTGGGTTGCCACTTTA | 127 | 59.8 |
| Murine hepatitis virus A59 | CGATGATTAAAGGCCCACAAC | CGTCTTTACGCACAGCAAAC | 99 | 59.8 |
| M13 phage | CTTTAACTCCCTGCAAGCCT | TCTTAAACAGCTTGATACCGATAGT | 101 | 59.8 |
| PBCV-1 | ATTGAGACTGGACTGGGAAAG | CAGAACGCGATGGAATGTTTAG | 574 | 63.7 |
| φKZ | CGGCAGGATCGATGTTCGTA | TCCAGGCTAGTACCCAGACC | 255 | 67 |
| φ6 L segment | CGATCAACACTGTCGTCATCAA | GCAGAACCGAAGGACGAATATC | 105 | 67 |
| Qβ | CGACCGTGGCCTATCTATAATG | CGCGGAACGTGGTATAACTAA | 127 | 58 |
| T4 phage | AAGCGAAAGAAGTCGGTGAA | CGCTGTCATAGCAGCTTCAG | 163 | 63.7 |
| Vaccinia virus | CGGCTAAGAGTTGCACATCCA | CTCTGCTCCATTTAGTACCGATTCT | 71 | 67 |

**SI Table 3: Primer sequences used for virus-specific qPCR assays.** Forward and reverse primer sequences, annealing temperatures, and expected amplicon sizes are shown for each virus-specific qPCR assay.

**SI Table 1 is attached as a .xlsx file**

**Author Contributions:**

NWW:

- Major contributor to framing, experimental design and writing of manuscript.
- Major contributor to selection and coordination to obtain viral mock community, generation of manuscript claims, figure design
- Performed VLP enrichment methods & viral stock extractions
- Performed Illumina and PacBio library preparation on Mock DNA 4-6 & Mock RNA 4-6
- Processed human intestinal biopsies with viralMEM & extraction.
- Major contributor providing feedback on analysis and interpretation of results.
- Contributed to bioinformatic processing of Mock DNA 4-6 and Mock RNA 4-6 samples post sequencing
- Performed assembly and viral binning of Mock DNA 4-6 and Mock RNA 4-6
- Analyzed and generated figures 1C, 1F, 2E-G, 3, 4C-E, SI Fig. 1, & SI Fig. 7-9
- Performed bioinformatics analysis of human Anellovirus
- Primary contributor to drafting the manuscript. Contributed to writing methods.
- Organized authors and defined deliverables during manuscript preparation.

AER:

- Major contributor to selection and coordination to obtain viral mock community, generation of manuscript claims, figure design
- Identified optimal low biomass workflows and handling steps
- Performed whole genome amplification on all samples
- Performed VLP enrichment methods & viral stock extractions
- Performed Illumina and PacBio library preparation on Mock DNA 1-2 & human intestinal biopsies
- Contributed to bioinformatic processing of Mock DNA 4-6 and Mock RNA 4-6 samples post sequencing
- Processed human intestinal biopsies with viralMEM & extraction
- Contributed to bioinformatic processing of Illumina sequencing of Mock DNA 1-2, 4-6, and Mock RNA 4-6
- Analyzed and generated figures SI Fig. 3-5, & SI Fig. 7
- Contributed to writing methods

CRN:

- Contributed to selecting and obtaining viral mock communities
- Performed phage propagation for T4, Qbeta, and PhiKZ
- Analyzed and generated figures 1A-B, 1D-E, 2A-D, 4B, SI Fig. 1, SI Fig. 2, SI Fig. 6, & SI Fig. 9
- Contributed to figure design
- Contributed to bioinformatic processing of Mock RNA 4-6 samples post sequencing
- Contributed to writing methods
- Performed phage bioinformatics analysis on human samples

AG:

- Performed viral specific qPCR's on pre and post PTA samples
- Performed phage propagation of PhiKZ
- Processed human intestinal biopsies with viralMEM & extraction
- Created Figure 4 panel A
- Contributed to writing methods

XPP:

- Contributed to development of low biomass practices utilized throughout the project
- Contributed to identification of low biomass extraction kit compatible with human clinical sample
- Contributed to initial pilot experiments characterizing PTA performance and background
- Identified and validated initial pilot on RT enzyme and clean-up method for Mock RNA community
- Developed and validated viral-specific qPCR workflows

ACB, JAM, and SSK:

- Recruited the participants, performed endoscopies to obtain biopsies, and edited the manuscript
- Obtained IRB approval

EVM:

- Edited manuscript and coordinated initial processing and shipments of the duodenal biopsies

RSC:

- Edited manuscript and participated in the design of the clinical study

BJ:

- Obtained funding, oversaw the whole clinical study, and provided leadership to the entire clinical team.
- Revised the manuscript

RFI:

- Provided funding, feedback on study design, technical guidance, manuscript feedback and editing, and leadership throughout the project

**Author Contact Information and ORCiDs:**

Eric V. Marietta:



Mock DNA (4–6) + NTCs
